# GPX4-VIM equates a proliferating DTP state in TNBC subtypes with converged vulnerabilities to autophagy and glutathione inhibition

**DOI:** 10.1101/2023.05.18.541287

**Authors:** Nazia Chaudhary, Bhagya Shree Choudhary, Sushmita Patra, Shivani Malvankar, Anusha Shivashankar, Eeshrita Jog, Vaishali V. Kailje, Sonal Khanna, Subhakankha Manna, Sarthak Sahoo, Soundharya R, Mohit Kumar Jolly, Sorab N. Dalal, Nandini Verma

## Abstract

Frequent metastatic relapses in Triple-Negative Breast Cancer (TNBC) patients with residual disease is a clinical challenge, largely due to tumor heterogeneity and absence of strategies that target proliferating chemo-tolerant cells. Here, we longitudinally modeled cellular state transitions from dormant drug-tolerant persister (DTP) into proliferating drug-tolerant persister (PDTP) in cells representing all TNBC subtypes. Combining subcellular imaging with phenotypic and biochemical assays, we identified distinct and converged spectrums of alterations in TNBC-PDTPs. We show that PDTPs retain acquired resistance with increased invasion potential. Moreover, Basal-Like DTPs enter into a non-reversible mesenchymal state while luminal androgen receptor-positive gain partial-Epithelial-to-Mesenchymal Transition (EMT) with vimentin upregulation. PDTP state dwells on high autophagy with reduced glutathione and GPX4 levels, rendering it vulnerable to autophagy suppression and ferroptosis. Interestingly, we find that GPX4 negatively regulates EMT and autophagy in TNBC, and an inverse correlation of GPX4-VIM expression along with autophagy genes predicts survival in TNBC patients undergoing chemotherapy.

## INTRODUCTION

Triple negative breast cancer (TNBC) is the most aggressive and heterogeneous type of breast malignancy with four distinct molecular subtypes that show diverse clinical behaviors and responses to therapy (*1*). Classification of TNBC tumor subtypes is primarily done based on their distinct transcriptional states but it also delineates the cell type of their origin (*2*, *3*). Basal-Like subtypes (BL-1 and BL-2) tumors are more epithelial in nature, while Mesenchymal Stem-Like (MSL) tumors are metastatic and are enriched in tumor stem cells. The Luminal Androgen-Receptor positive (LAR), a minor fraction of TNBC tumors that are luminal in origin that express Androgen receptors (*2*). As compared to non-TNBC, TNBC patients show the worse prognosis and have shorter disease-free and overall survival rates, mainly due to disease relapse with resistance to therapy and metastasis (*1*, *4*, *5*). Tumors under different TNBC subtypes show varied responses to therapy, as BL-2 subtype tumors are more chemotherapy-refractory in nature, while LAR subtype has the lowest pathologic complete response (pCR) rate and exhibits a higher degree of drug resistance (*2*, *6*, *7*).

Conventionally, clinical management of TNBC relies on chemotherapeutic agents as first-line therapy, mostly in a neoadjuvant setting due to the lack of druggable hormone receptors (estrogen receptor, ER and progesterone receptor, PR), or human epidermal growth factor receptor-2 (HER2) overexpression (*8–10*). TNBC shows a better response to chemotherapeutic agents owing to a higher proliferation index than non-TNBC. It is worth recognizing that a good proportion of patients achieve initial pCR (*11*, *12*), however, unfortunately a substantial fraction of patients shows early relapse within 2-5 years post-therapy with distant metastases (*13*). Moreover, a large proportion of TNBC patients (∼50%) do not reach pCR and have post-therapy residual disease with metastasis (*9*, *11*). Survival of TNBC patients with chemo-tolerant residual disease as well as with relapsed chemoresistant disease is often extremely poor (*11*, *13*, *14*). Clinical management of TNBC patients that do not achieve pCR with first-line chemotherapeutics is very challenging due to a very limited understanding on the nature of this residual disease and targeted therapy options for this group (*5*, *14*, *15*). Therefore, there is a dire need to understand molecular and cellular characteristics of chemo-tolerance in different TNBC subtypes that can pave the way for more precise treatment options.

Growing number of evidences from different cancer types such as non-small lung cancer, melanoma, glioblastoma, colorectal, and breast cancers support the ubiquitous idea that drug-tolerance against targeted therapies and cytotoxic agents in solid tumors is acquired mainly through epi-genetic mechanisms or alteration in cellular metabolism (*16*, *17*). Drug-induced molecular re-programming events in cancer cells result in a reversible drug-tolerant cellular state that can persist under high doses of therapy (*18*, *19*). These drug-tolerance-associated adaptive cellular mechanisms are largely known to support cell survival, however, in parallel often confer distinct changes in cell phenotypic states and susceptibilities or molecular/pathway addictions (*18*, *20–23*). For example, recent studies across different cancer types indicated that activation of the EMT program led to inactivation of the apoptotic machinery and the acquisition of multi-drug resistance (*24*, *25*). The drug-tolerant persister (DTP) cancer cells that can withstand high doses of therapeutic agents are shown to have altered cellular metabolism to support acquired changes in cellular plasticity (*19*, *24*, *26*). One of these metabolic shifts in DTP cells is increased levels of polyunsaturated fatty acids (PUFA) synthesis, though, it also imparts susceptibility to iron-dependent ferroptotic cell death owing to increased membrane lipid peroxidation (*27–29*). More importantly, DTP cancer cells from different cancer types are found to have vulnerable redox management due to significant suppression of proteins like glutathione peroxidase 4 (GPX4) that detoxifies peroxidated lipids and thereby prevents the damage of cell membranes lipids and inhibit ferroptosis (*18*, *29*, *30*). Studies in lung and colorectal cancers have shown that this unique phenotype of DTP cells provides a tangible therapeutic opportunity to eliminate apoptosis-resistant cancer cells by inducing ferroptotic cell death by inhibiting GPX4 or system Xc^-^ (*30–32*).

Triple-negative breast tumors are highly proliferative and metastatic in nature, and show an enhanced susceptibility to pharmacologic inhibition of glutathione biosynthesis pathways (*33*). Moreover, acyl-CoA synthetase long-chain family member 4 (ACSL4) that facilitates enrichment of PUFA in the cell membrane is shown to be preferentially upregulated in Basal-Like breast cancer cells and its expression correlates to sensitivity to ferroptosis (*34*). In addition to this, our previous study demonstrated that TNBC cells are intrinsically susceptible to ferroptosis-inducing agents like Fin56, Erastin and BRD4 targeted therapies that can induce intra-cellular reactive iron and GPX4 downregulation in TNBC cells and mice xenograft (*35*). Moreover, we reported that human TNBC tumors differentially express genes involved in iron homeostasis, glutathione metabolism and reactive oxygen species (ROS) regulation generating a ferroptosis-promoting gene signature. Additionally, TNBC tumors has a remarkably lower gene and protein expression levels of key anti-oxidants: GPX4, GSS (Glutathione synthetase), and iron transporter protein ferroportin (Fpn-1/SLC40A1), while elevated levels of iron binding receptor, FTH1 and ACSL4 (*35*). Since cellular sensitivity towards ferroptosis can be exploited to target drug-tolerant cancer cells, it will be crucial to investigate whether TNBC cells belonging to different molecular subtypes enter to a chemotherapy-tolerant state with a similar or differential status of vulnerability to ferroptosis inducers.

Here we have longitudinally modeled the development of proliferating drug-tolerant persister (PDTP) cells from a latent DTP cellular state in cell lines representing different TNBC subtypes in response to different classes of chemotherapeutics. We find that cells from all four TNBC subtypes were capable of developing DTP cell populations and after a quiescent phase entered a proliferative cycle and showed increased resistance to chemotherapeutics when compared to the parental cells (PC). PDTPs from Basal-Like and LAR subtypes gained varied degrees of Epithelial-to Mesenchymal transition (EMT) states with an increase in vimentin levels. Transmission Electron Microscopy (TEM), gene and protein expression analysis with cellular assays revealed that TNBC PDTP cells display some common alterations in sub-cellular organelles and biochemical composition of the PDTP cells that results in elevated lipid reactive species, suppressed glutathione synthesis pathway and higher basal autophagy that render them susceptible to ferroptosis inducers and autophagy inhibition, respectively. Most importantly, results from GPX4 inhibitor, RSL3-tolerant TNBC cells and lentivirus-mediated GPX4 expression modulation affirm that GPX4 regulates the expression of mesenchymal genes and autophagy in TNBC. Immunofluorescence (IF) staining in human breast cancer tissues coupled with multiple in-silico analyses of breast cancer patient datasets, identify a reverse correlation between GPX4 and vimentin that, together with autophagy markers can strongly predict survival and response in chemotherapy in TNBC patients with aggressive disease.

## RESULTS

### Chemotherapy induces a DTP state in all TNBC subtypes that progresses to a PDTP state with acquired drug resistance and higher proliferation

To study tolerance to chemotherapeutic agents in TNBC we developed cellular models of paclitaxel-, doxorubicin- and cisplatin-tolerant persister cells from cell lines representing each subtype of TNBC as described (*18*, *21*, *36*) (Fig. 1 and Fig. S1). We generated dose response curves to establish the IC50 for each drug (Fig. 1SA-C) for the chemotherapeutic drugs in the TNBC cell lines (hereafter referred to as parental cells (PC)) MDA-MB-468 (BL-1 subtype), HCC70 (BL-2 subtype), MDA-MB-231 and HS578T (MSL subtype) and MDA-MB-453 (LAR subtype). We chose chemotherapy doses that result in the survival of up to 15% of the cell subpopulation post-treatment in 72-96 h (Fig. 1A). These drug-tolerant TNBC residual cells endured an additional cycle of therapy without cell death, a subpopulation that corresponds to the minimal residual disease in human tumors (37–39). It was observed that the TNBC DTP cells could tolerate 50-250 folds of IC_50_ concentration of paclitaxel (Fig. 1E) while the maximal tolerated doses for doxorubicin and cisplatin were relatively lower at, 4.3 and 20.4 folds of IC_50_, respectively (Fig. S1F).

**Figure 1:**
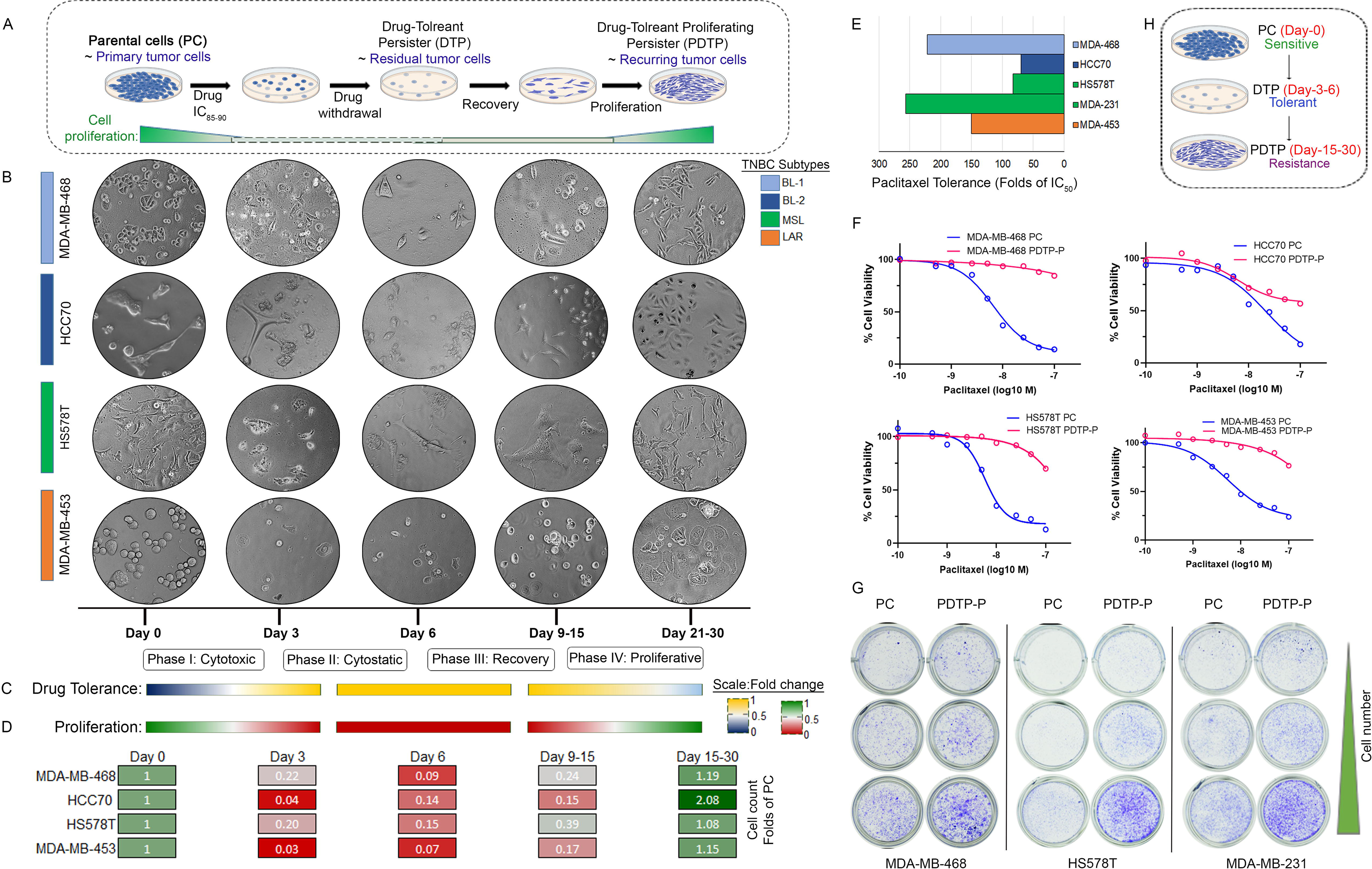
Longitudinal modelling of drug-tolerance to generate proliferating persisters, PDTP cells in different TNBC subtypes. (A) Schematic representation of process of developing TNBC PDTP from dormant DTP cells generated by chemotherapy treatment of parental cells, PC. TNBC cell line representatives from different molecular subtypes MDAMB-468, HCC70, HS578T and MDA-MB-453, were treated short term with chemotherapy (IC_80-95_) cytotoxic doses in vitro. Proliferation of dormant residual tumor cells was observed after a long-term drug holiday. (B) Phase contrast micrographs (20x magnification) of the TNBC cell lines in different time defined longitudinal cellular phases during the development of proliferating drug-tolerant persister from each subtype (color bars in left represent subtypes: light blue-BL-1, dark blue- BL-2, green-MSL and orange-LAR) using paclitaxel (0.5-1μM) treatment as a representative chemotherapy agent. (C) Drug tolerance represented as fold was calculated by percent of cells surviving the persister deriving doses at each phase assuming untreated PC with no tolerance, (color key, blue to yellow depicts 0-100% tolerance). (D) Proliferation rates of TNBC cell lines under different phases of development of PDTPs calculated by counting the cells in each time points shown, and represented as folds of parental cells, where the value is represented as 1 (color key, green to red depicts cell no from 100 to 0 %). (E) Tolerance to paclitaxel in different TNBC cell lines represented ad Folds of paclitaxel IC_50_ in TNBC cell lines. (F) Dose-response curves of varying concentrations paclitaxel in the PC and Paclitaxel PDTP (PDTP-P) derived from indicated cell lines (MDAMB-468, HCC70, HS478T and MDA-MB-453) treated for 72 hours. Dose-response curves are presented as means of three replicates. (G) Colony seeding potential detected by colony formation assay in PC and PDTP-P of respective TNBC cell lines. (H) Schematic representation summarizing development and characterization of DTP and PDTP from PC shown in figure 1.

After drug withdrawal, we have longitudinally followed and monitored the post-therapy residual drug-tolerant cells to observe the time-dependent changes in morphological features, drug tolerance, and cell proliferation for 30 days (Fig. 1B-D). We have observed that cells from all the TNBC subtypes underwent an initial cytotoxic phase (Phase I) in the first 72h post-paclitaxel treatment, where accelerated cell death was observed in the presence of drug (Fig. 1B, D). After the withdrawal of drug we observed a cytostatic phase (Phase II) in which remaining residual DTP cells showed no significant changes in cell numbers for the next 3 days and remained quiescent. The quiescent cells entered into a slow-dividing phase (Phase III) within 15 days of post-treatment and the cells started to gain mitotic recovery (Fig. 1B,D). Eventually, it was observed that the DTPs entered in a proliferation phase (Phase IV) having cell division rates equivalent to greater than the PC, as in case of HCC70 recovered DTP cells, these were more than 2 folds proliferative than PC. In phase IV cells were stably passaged as proliferating DTP (PDTP) used for further characterization (Fig. 1A). Importantly, we were successful in generating PDTP cells not only with paclitaxel (PDTP-P) but also with doxorubicin (PDTP-D) and cisplatin (PDTP-C) from all the four TNBC subtypes (Fig. S1E) using similar treatment regimes.

Phase contrast microscopy analysis of the DTP cells under phase II and III suggested striking changes in the morphological features of the cells (Fig. 1B), especially, lines corresponding to Basal-Like subtypes (MDA-MB-468 and HCC70) that were stable even in PDTPs (Fig. 1B, Fig. S1E). We have observed that irrespective of the therapy agent used, acquisition of drug tolerance in MDA-MB-468 and HCC70 was coupled with loss of cell-cell contacts and gain of spindle shaped cellular morphology, characteristic features of a mesenchymal state (*22*, *24*, *37*). TNBC cell lines from MSL subtype, HS578T and MDA-MB-231 did not show an evident change in morphology; however, notably, we observed some transient loss of cell-cell contacts and morphological alterations in LAR cells, MDA-MB-453 in DTP state that partially retrogressed in the PDTP state with the acquisition of cell-cell contacts (Fig. 1B).

We then sought to determine whether the PDTP cell population retains the same level of drug tolerance as the dormant or slow-dividing DTP cells by performing dose-response curves for PC and PDTP-P cells in each TNBC subtype (Fig. 1F). It was observed that the entire PDTP-P population was not equally tolerant to higher doses of paclitaxel and there was a significant amount of cell death with dose escalation. However, the PDTP-Ps were more resistant to paclitaxel as compared to the PCs. These results suggest a decrease in the drug tolerance of PDTPs after gaining proliferation in the absence of the drug, suggestive of phenotypic reversibility owing to a mechanism that is mostly non-genetic in nature (*36*, *38*). Several previous studies support the idea that heterogeneous cell population could contribute to distinct clonal evolutionary lineages of epigenetic re-programing in the drug-tolerant cells (*37*, *39–41*). Further, as drug-resistant cells show an increased ability to form tumor across tumor types including TNBC tumors (*13*, *14*, *42*) we tested the colony formation potential of TNBC PDTP-P cells (Fig. 1G). As seen in the results of colony formation assays (Fig. 1G). Using different cell seeding densities of PDTP-Ps in different TNBC subtypes we found that they could form more viable colonies (even at low density) compared to their corresponding PCs. Together these results underline a substantial capacity of TNBC cells, irrespective of subtype, to attain a DTP state with diverse classes of chemotherapy agents and to overcome the proliferation pause in the absence of drug treatment (Fig. 1 H). These characteristic behaviors of drug-tolerant cell models can explain the basis of early and aggressive disease relapse observed for post-treatment TNBC tumors in the clinic (*13*).

### Chemotherapy induces a non-reversible mesenchymal state in BL-1 and BL-2, while a partial-EMT is LAR subtype PDTP cells

As drug-induced molecular reprogramming often results EMT in epithelial tumors, a plastic state that is more drug-resistant in nature (*24*). We observed atypical changes in morphological features in chemotherapy-tolerant DTP and PDTP cells derived from Basal-Like TNBC cell lines (MDA-MB-468, BL-1; and HCC-70, BL2) that are indicative of induction of EMT program (Fig. 1B. and S1E). To study the cell-cell contact and epithelial geometry, we performed TEM analysis of PC and PDTP monolayer cells (Fig. 2A). As seen in the TEM images, MDA-MB-468 and HCC70 PC show an epithelial morphology with apicobasal cell polarity and well-established cell-to-cell contacts (magnified inserts) which were totally lost in their respective proliferating persisters drove from paclitaxel (PDTP-P), doxorubicin persister (PDTP-D) and cisplatin persister (PDTP-C) (Fig. 2A). Moreover, PDTP cells have obtained an elongated morphology with front/back polarity, which can be accredited to cytoskeleton-driven protrusion formation, indicative of higher cell motility phenotype. In order to verify that these morphological changes are indeed due to EMT, we have analyzed the expression and localization of key EMT markers. Immunofluorescence staining of E-cadherin and vimentin in MDA-MB-468 (Fig. 2B) and HCC70 (Fig. 2D) show a significant decrease in E-cadherin expression and its delocalization from cell-cell contacts, while there is a dramatic increase in expression of vimentin (Fig. 2C, D) in all PDTP cells. More importantly, these changes were transcriptionally controlled, as the gene expression analysis of these EMT-markers showed significant alterations in expression in the PDTP cells (Fig. 2F). It was observed that the mRNA level of vimentin was increased in Basal-Like cells MDA-MB-468, MSL subtype cells MDA-MB-231 and HS578T (Fig. 2F). A similar increase was observed when the protein levels of vimentin and E-cadherin were measured in Western blot assays (Fig. 2G) showing a strong upregulation (upto 27 folds) of vimentin and loss of E-cadherin in the BL-1 and BL-2 PDTP cells (Fig. G, H). However, interestingly, PDTP derived from LAR cell line MDA-MB-453, showed an increase in the levels of both vimentin and E-cadherin suggesting that these cells have a hybrid epithelial and mesenchymal state (*6*, *43*). Recent research has advocated the role of partial EMT or hybrid EMT in metastasis and secondary tumor formation in breast and other epithelial type of tumors, as these cells have a higher potential to revert back to an E-cad^+^ epithelial cell state while most of the full EMT cancer cannot overcome the mesenchymal state (*44–49*). These results indicate that chemotherapeutics induced the development of drug-tolerant persister cells that are phenotypically distinct from their parental cells and are enriched in mesenchymal phenotype with a ubiquitous increase in vimentin in all TNBC subtypes (Fig. 2I).

**Figure 2:**
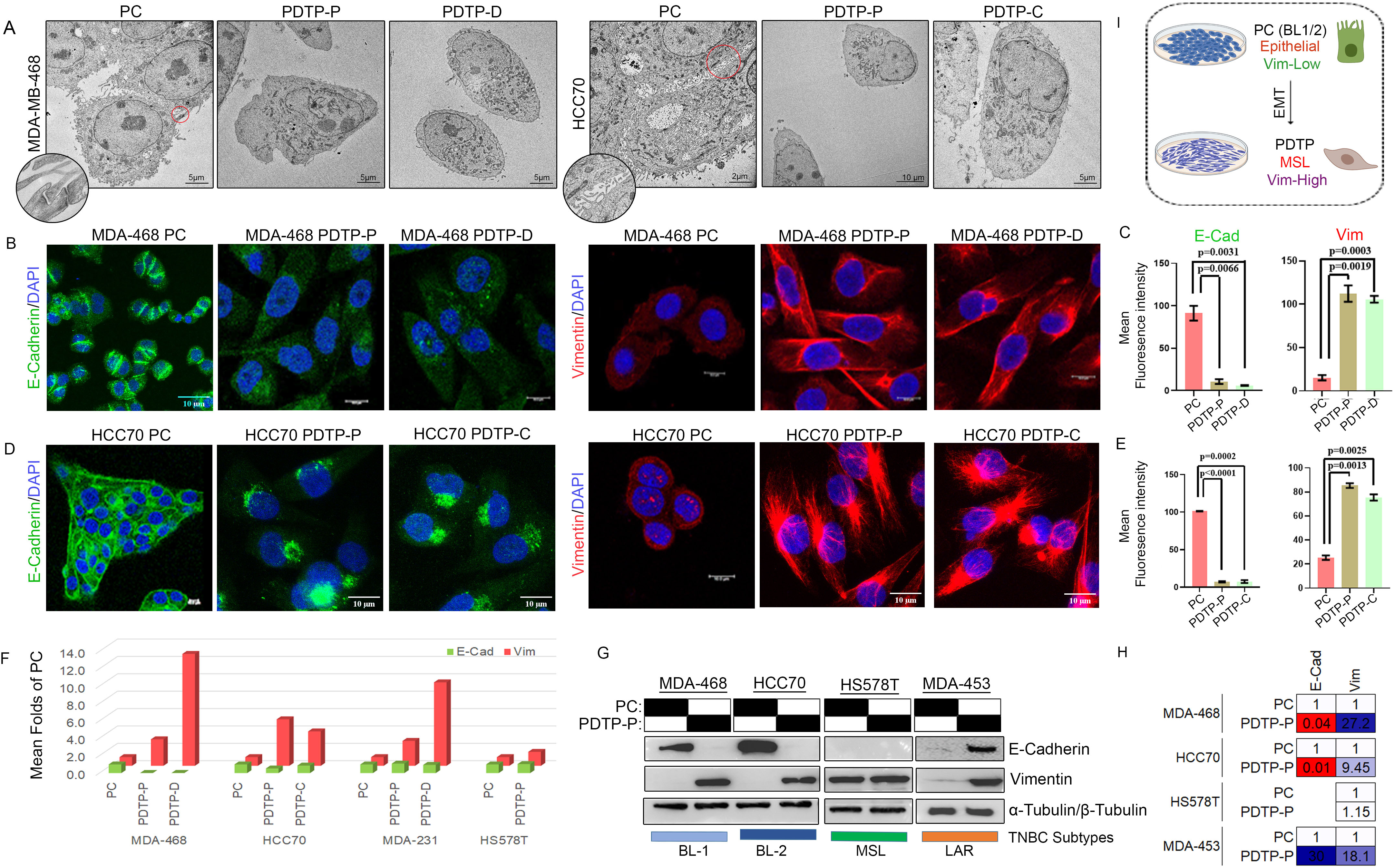
Proliferating chemotherapy-tolerant persisters from Basal-Like subtype attained irreversible EMT phenotype while LAR acquired a partial EMT state. (A) TEM images of intercellular spaces revealed loss of cell-cell contact in PDTP cells and intact tight junctions between cells in PC (indicated in zoomed inserts). Scale bars, 5 or 10μm, as indicated. (B, D) Representative confocal microscopy images of PC and PDTP cells after immunofluorescence staining for analysing the expression and localization of E-cadherin and vimentin. Green fluorescence indicates E-cadherin and Red fluorescence indicates vimentin. Scale bar: 10 μm. (C, E) Bar graphs representing the quantification of mean fluorescent intensity of PC and DTTP for E-cadherin and Vimentin immunofluorescence staining shown in B and D in with statistical significance of p < 0.001. (F) The expression levels of E-cadherin and vimentin mRNAs were assessed in different TNBC subtypes using QPCR analysis with gene specific primers and were normalised to actin expression. Gene expression in PDTP cells were plotted as mean fold change compared to PC. (G) Protein expression of E-cadherin and vimentin in PC and PDTP-P as assessed by Western bolt analysis in 4 TNBC subtypes, α-tubulin protein expression was used as loading control (H) ImageJ software was used for quantification of Western blot band intensities of PC and PDTP-P in G and presented as heat map of changes folds of PC. (I) Schematic representation summarizing main results of phenotypic changes in PDTP TNBC cells shown in figure 2.

### PDTP cells exhibit cytoskeletal reorganization with enhanced migration and invasive potentials

Drug resistant residual disease in patients with TNBC is often coupled with distant metastasis and poor survival (*14*, *50*, *51*). As we have observed dramatic cellular changes and EMT alterations in PDTP cells (Fig. 2), we sought to further analyze the changes in their cytoskeleton and capability for migration and invasion, which are associated with a high mesenchymal states of persister cells (*24*, *25*, *52*). Immunofluorescence staining and confocal microscopy imaging of filamentous actin fibers in both BL-1/2 subtypes (Fig. 3A) and in MSL (Fig. S2) TNBC cells indicates a significant increase in actin-based cytoskeleton (Fig. 3B) and actin polymerization in PDTPs driven by paclitaxel, doxorubicin and cisplatin. These cytoskeletal changes indicate a gain of migratory capability of the PDTPs (*53*, *54*). Further, wound healing assays demonstrated that the PDTP cells showed an increased migration rate and the wound was closed to a greater degree (Fig. 3C-E). As seen in the figure, MDA-MB-468 PC which is minimally migratory in behavior, PDTP showed a dramatic increase in cell migration. In case of MSL cell lines that are usually migratory in nature, there is an enhancement in migration and wound healing potential (Fig. 3D, E) of PDTP cells. Next, we analyzed the changes in metastatic potential using Matrigel invasion assays in PC and PDTP-Ps (Fig. 3F, G). We have observed that paclitaxel-PDTP cells gained up to 5 folds increase in the invasion capacities (Fig. 3G) as compared to PCs, and these changes were seen in all three subtypes of cells tested. These data suggest that chemo-tolerant TNBC persister cells are not only phenotypically enriched in mesenchymal properties but also have functional capacities for a higher migration and invasion compared to drug-sensitive parental cells (Fig. 3H). These findings are in line with the known metastatic potentials of drug resistant cancer cells (*55*, *56*) that make the disease more aggressive and challenging to treat.

**Figure 3:**
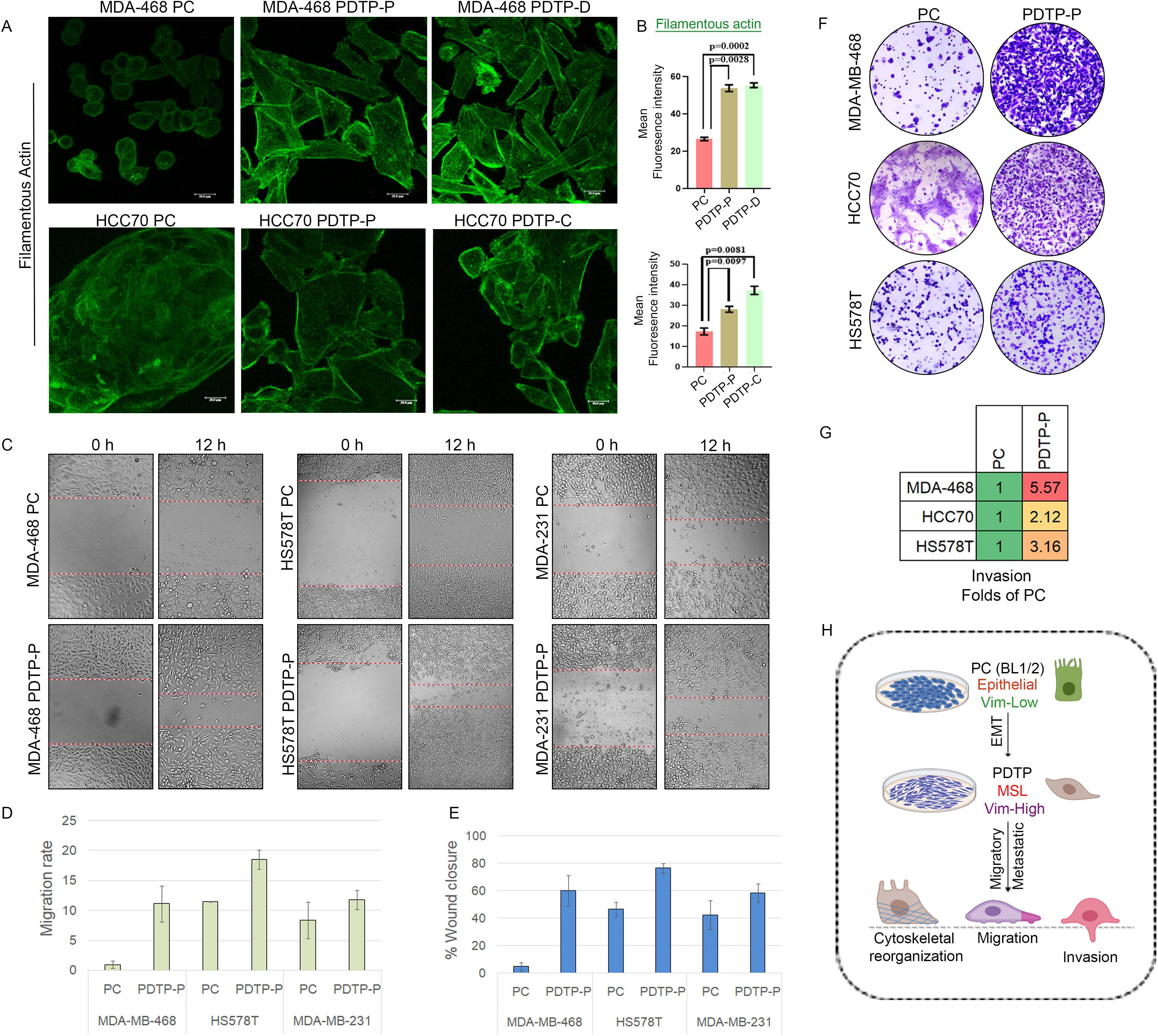
TNBC PDTP cells show increased migration and invasion potential with massive cytoskeletal reorganization. (A) Representative confocal microscopy images of immunofluorescence staining of filamentous actin in Basal-Like TNBC PC and their respective PDTP cells. (B) Mean fluorescent intensity of filamentous actin was estimated from the replicates PC and their PDTP-P, PDTP-C and PDTP-D and plotted in the bar graphs with statistical significance of p < 0.001. (C) Bright field microscopy images (20x magnification) of cell migration through scratch wound healing assays in TNBC PC cells and PDTP-P were recorded at 0 and 12 hours. Red dotted line indicates the migration front of the cells (D) Bar graph representing the migration rate and (E) percentage of wound closure in PC and PDTP-P TNBC cells calculated as indicated in the material and method section. (F) Bright field microscopy images showing transwell matrigel invasion assay in PDTP-P and PC TNBC cells. (G) Fold difference in invasion capacity compared to PC is presented in heatmap. (H) Schematic representation to summarize the increased cellular motility and invasive behaviour in TNBC PDTP cells as shown in figure 3.

### PDTP cells have elevated autophagy flux and lysosomal content that render them susceptible to autophagy inhibitors

EMT changes in response to high-dose drug assault not only creates a stressful environmental and intrinsic conditions that a cancer cell has to survive, but also makes a sudden shift in the amount of energy required by the cell to handle enhanced cytoskeletal reorganization to support cell migration and invasion (*24*, *57*). A growing number of studies demonstrate that cellular stress conditions induce autophagic flux and higher lysosomal activity that assist the cancer cells to cope with increased energy requirements even in cells under EMT state (*58–61*). Very interestingly, the Transmission Electron Microscopy studies of TNBC PDTP cells across the subtypes showed a striking increase in the autophagosomes and lysosomes in comparison to the respective PCs (Fig. 4A). As seen in the TEM images, PDTPs derived from all the chemotherapeutic agents were able to induce enhanced autophagosomes (solid red arrow) indicated by membrane bound vacuolated subcellular structures and lysosomes (solid yellow arrow) seen as electron dense structures in the proliferating persister cells. In order to confirm that the subcellular changes seen in the TEM are indeed associated with induction of autophagy, we performed IF staining and confocal microscopy analysis using specific molecular markers for autophagy, LC3B (Fig. 4 B) and for lysosomes, LAMP1 (Fig. 4C) in different PC and PDTP cells. Results of these IF studies clearly indicate that there is an extensive increase in LC3B positive puncta (Fig. 4B) in PDTPs indicating higher autophagy flux and higher lysosomal content indicated by large LAMP1 positive perinuclear vacuoles (Fig. 4C). Further, we performed western blot analysis to probe for autophagy markers LC3B lipidation and P62 proteins in PC and PDTP-P (Fig. 4D), and the results verified the increase in LC3B expression and its lipidated form.

**Figure 4:**
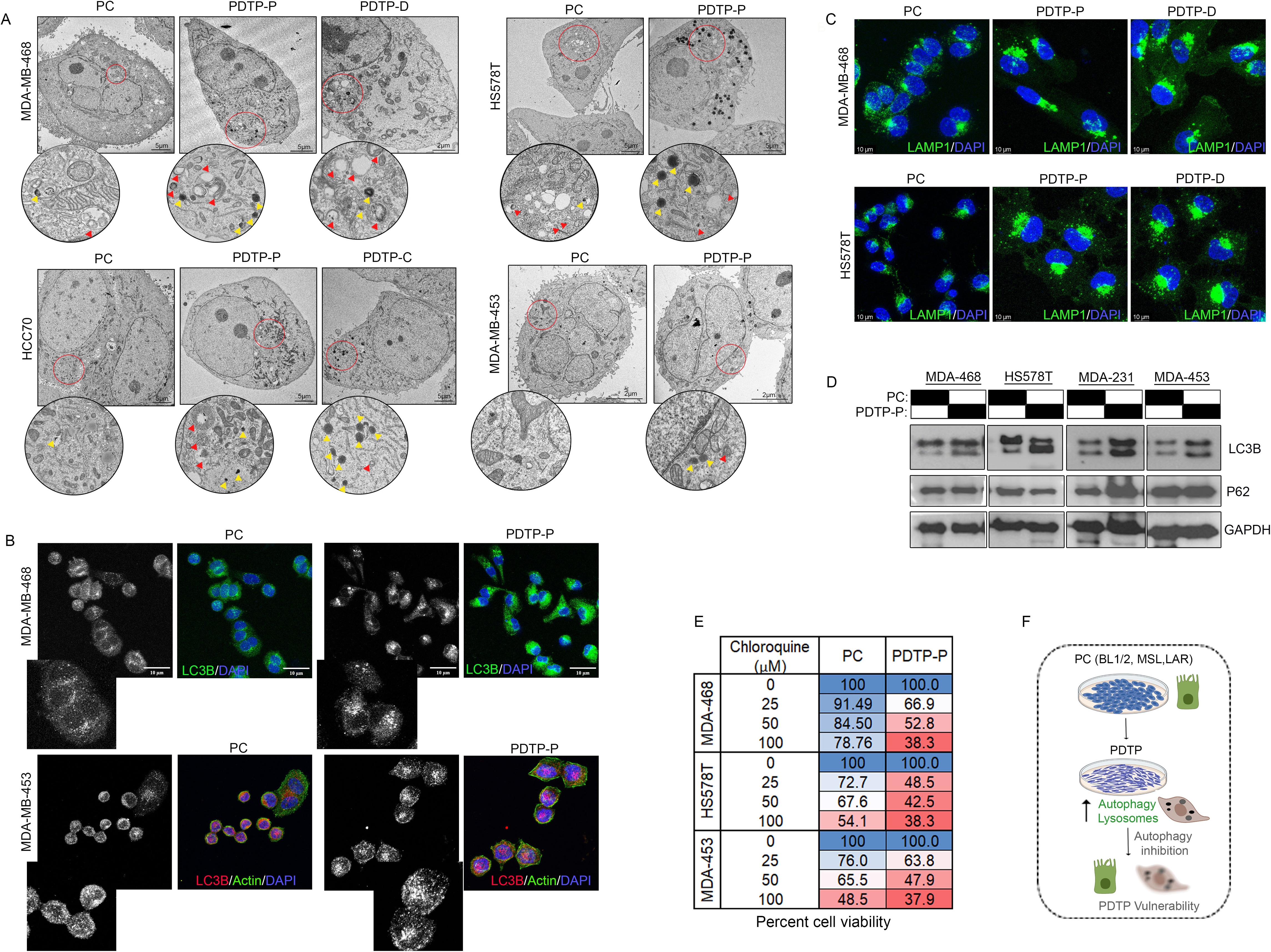
Enhanced autophagy in TNBC PDTP persister is a targetable vulnerability. (A) Representative TEM micrographs and zoomed inserts of red encircled areas showing increased number of autophagosomes (red arrow head) and lysosomes (yellow arrow head) in PDTP-P, PDTP-D and PDTP-C as compared to PC. Scale bar: 2 to 5 µm, as indicated. (B) Representative confocal micrographs showing immunofluorescence staining of autophagy marker LC3B and (C) lysosomal marker LAMP1 in PC and PDTP-P, PDTP-D in different TNBC cell lines. Scale bar: 10 µm. (D) Western blot analysis of autophagy related proteins LC3B and p62 in parental TNBC cell lines and their PDTP-P. GAPDH served as a loading control. (E) MTT assay was performed to check the viability of parental TNBC cells and PDTP-P treated with different concentrations of autophagy inhibitor chloroquine as indicated, data presented as mean percent cell viability in the heatmap. (F) Schematic representation to summarize data shown in figure 4 that suggests increased autophagy in TNBC PDTP renders them vulnerable to chloroquine.

To evaluate whether PDTP cells are using autophagy as a source of energy for their survival and maintenance, we have tested the sensitivity of these cells to increasing doses of autophagy inhibitor, chloroquine using the MTT assays (Fig. 4E). Percent cell viability (Fig. 4E) shown in the heatmap indicates that PDTP-P cells from different TNBC subtypes were significantly more sensitive to autophagy inhibition that indicated the survival of PDTP cells is considerably dependent on autophagy. These results collectively suggest that there is an increase in autophagosomes and lysosomal content in PDTP cells that support their cell survival, and highlights a unique vulnerability of PDTPs to autophagy inhibitor chloroquine (Fig. 4F) that can have therapeutic implications (*23*, *62*).

### PDTP cells exhibit elevated reactive iron burden with compromised GPX4-mediated anti-antioxidant defense mediating a higher vulnerability to ferroptosis inducers

Several recent studies have suggested that therapy-tolerant persister cells in epithelial tumors are deficient in anti-oxidant systems, including GPX proteins and are vulnerable to GPX4 inhibition (*18*, *20*, *63*). One of the critical features of oxidative damage at the sub-cellular level is aberrant mitochondrial structure and function (*64*). Moreover, it is also seen that the redox state of DTP cells can also be regulated by mitophagy (*65*, *66*). In order to examine the oxidative status and its associated damage to mitochondria in different TNBC PDTPs, we analyzed mitochondria integrity and morphological features using TEM (Fig. 5A). Electron micrographs of PDTP cells reveal anomalous alterations in the mitochondrial architecture, such as dense and shrunken mitochondria with frequently ruptured or missing mitochondrial cristae, all pointing toward a typical morphological characteristic of oxidative damage and peroxidation of membrane lipids (*64*). These features in all PDTP TNBC cell lines were seen irrespective of chemotherapy agent, indicating a similar mitochondrial phenotype drives drug-tolerance in TNBC. To understand the dynamics of redox homeostasis in PDTP cells we then estimated total ROS and lipid peroxidation using cell DCFDA assays and dual fluorescence BODIPY 581/591 C11 lipid probes, respectively, upon treatment of GPX4 inhibitor RSL3 or Erastin. As seen in the figure 5B, RSL3 induced higher cellular ROS in all the PDTP cells tested, the highest (upto 5 folds of PC) effects were seen in MDA-MB-231 and MDA-MB-468 cells. Similarly, a sharp red-to-green shift in the fluorescence of the BODIPY 581/591 C11 lipid probes in PDTP cells upon Erastin treatment indicates very high lipid peroxidation in PDTP cells compared to PCs (Fig. 5C and S3A, B). These results suggest that drug-tolerant cells have a compromised anti-oxidant defense and are more dependent on glutathione than their parental counterparts which probably create an enhanced ferroptosis susceptibility (*33*, *35*). In order to investigate further to determine whether the higher ROS induction is indeed associated with lower levels of cellular glutathione levels, we estimated total cellular glutathione levels in PDTP cells in steady state and RSL3 treated conditions (Fig. 5D). Interestingly, it was observed that though the PDTP cells have up to 40% lesser cellular glutathione levels in untreated conditions in comparison to PCs, while the RSL3 treatment resulted in a severe drop (up to 80% compared to PCs treated with RSL3) in the cellular glutathione levels in PDTP cells (Fig. 5D).

**Figure 5:**
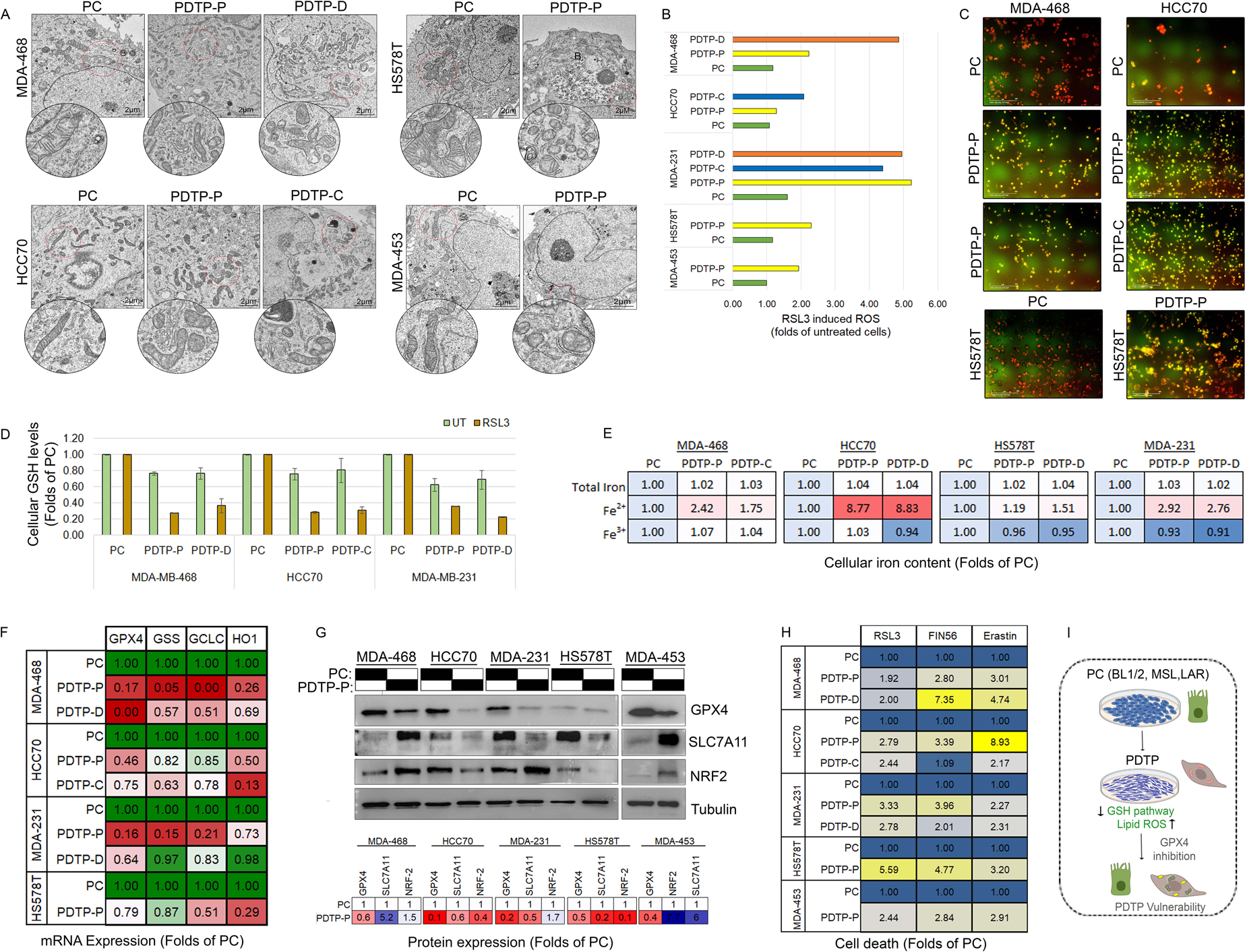
TNBC PDTPs are more sensitive to ferroptosis induction due to elevated mitochondrial damage, lipid ROS, labile iron coupled with glutathione depletion as a result of GPX4 pathway suppression. (A) Representative TEM micrographs and their zoomed inserts indicating change specific to mitochondrial structure in PDTP-P, PDTP-D, and PDTP-C as compared to PC in the indicated TNBC cell lines. Scale bar- 2µm, as indicated. (B) Cellular ROS levels were analysed by DCFDA assay in RSL3 treated parental TNBC cells and their PDTP-P, PDTP-C and PDTP-D, represented in the bar graph as folds of untreated cells as control. (C) Representative fluorescence microscopy images (20x magnification) acquired by Incucyte live-cell analysis system to detect lipid peroxidation after exposure to Erastin (1μM) for 4 hours followed by incubation with lipid probe BODIPY 581/591 in PC, PDTP-P and PDTP-C of indicated TNBC cell lines. A red to green shift in fluorescence show lipid peroxidation in PDTPs. (D) Parental TNBC cell lines and indicated PDTPs were treated with RSL3 (1μM, 4h) and cell lysates were used to determine cellular GSH levels using GSH-Glo Glutathione assay kit as described under method section. Total Cellular GSH levels were plotted folds of PC, mean±SD from 3 replicates of the assay (E) Cellular levels of ferrous, ferric and total iron were measured in PDTP-P, PDTP-D and PDTP-C in steady state condition in comparison to PC of indicated TNBC cell lines using iron assay kit. Results represent mean of 3 replicates. (F) The mRNA expression level of genes involved in ferroptosis regulation namely, GPX4, GSS, GCLC and HO1 were analysed using QPCR analysis in PDTP-P, PDTP-D and PDTP-C in comparison to PC of indicated TNBC cell lines, mean fold changes are represented as heatmap. (G) The protein expression levels of GPX4, SLC7A11 and NRF2 in parental TNBC cell lines and PDTP-P were analysed by WB analysis. ImageJ software was used to quantify the band intensities and fold change compared to PC from replicates is presented as heatmap. (H) MTT assay was performed to determine viability of indicated TNBC cell lines and PDTP-P and PDTP-D as compared to PC treated with RSL3, FIN56 and Erastin. Percent Cell death was calculated in each case and presented as folds of PC in the heatmap as mean from triplicate experiments (I) Schematic representation of the proposed model for GPX4, GSH inhibition and increase in lipid ROS resulting to ferroptosis vulnerability in PDTP cells is depicted.

Lipid peroxidation in cells is classically mediated by increased reactive iron pools in the cells that participate in the Fenton reaction and generate lipid ROS resulting in ferroptotic cell death (*67*), we next estimated the total cellular iron content and its reactive labile state (Fe^2+^) in steady state (Fig. 5E) and RSL3 treated (Fig. S3C) conditions in PDTPs and PCs. The results of cellular iron assays indicate that although the total content of intracellular iron remains largely unchanged, there is a substantial increase in the labile iron in all the PDTPs, HCC70 it was the highest (up to 8 folds of the PC) (Fig. 5E). Similarly, in RSL3 treated PDTP cells Fe^2+^ were up to 2 folds higher as compared to RSL3 treated PCs (Fig. S3C). These results with increased cellular ROS (Fig. 5B, C) and decreased total glutathione levels (Fig. 5D) indicate that the PDTP cells might have a transcriptional reprogramming that constantly support the high ROS cellular status. Hence, next we analysis of expression of GPX4 and several other key ferroptosis pathway molecules at both mRNA (Fig. 5F) and protein (Fig. 5G) levels using QPCR and WB analysis, respectively. The mRNA expression analysis shows a significant downregulation of GPX4 expression and other glutathione/ROS management pathway molecules, including GSS, GCLC, HO-1, xCT, and transcriptional regulator NRF-2 (Fig. 5F). Many of these changes were also observed at protein levels (Fig. 5G and S3D), specifically, GPX4 and HO-1 were consistently downregulated in PDTPs of almost all the subtypes as compared to the parental cells. Immunofluorescence staining of GPX4 in different PDTPs also showed a significant decrease in the cytosolic expression of GPX4 (Fig. S3E).

Because we have observed dramatic increased in cellular ROS and iron levels in PDTP, together with lower glutathione levels, we speculated that these cells might show minimal resistance to ferroptosis inducers (*31*, *67*). Therefore, we have tested the response of GPX4 inhibitors (RSL3 and FIN56) and xCT blocker Erastin for their potential cell death-inducing capacity in PDTPs derived using different chemotherapy (Fig. 5H). Results of cell viability assays indicate that TNBC PDTP cells are much more sensitive to inhibition of the xCT and GPX4 (Fig. 5H), which indicates a significant susceptibility to ferroptosis. These results together suggest a differential status of glutathione and NRF-2 derived anti-oxidant mechanisms in drug-tolerant persister TNBC cell lines that can be preferentially targeted by GPX4 inhibitors or ferroptosis inducing agents (Fig. 5I) and can create a specific therapeutic opportunity.

### RSL3 PDTPs reveals involvement of GPX4 in controlling mesenchymal phenotype and autophagy flux in TNBC

In the earlier study, we have demonstrated that TNBC are intrinsically vulnerable to ferroptosis and express lower levels of GPX4 and GSS, key ferroptosis pathway proteins as opposed to the non-TNBC counterparts (*35*). In light of the above findings (Fig. 5) that chemo-tolerant TNBC substantially downregulates GPX4 and increases labile iron pools that render them even extremely susceptible to glutathione pathway inhibitors, we hypothesize that PDTP phenotypes are at least partially dependent on GPX4 expression in TNBC. In order to investigate in this direction, we took an indigenous approach to develop RSL3 PDTPs from different TNBC subtypes with an idea to deplete a larger TNBC cell population that is susceptible to ferroptosis induction by GPX4 inhibition while enriching a very small fraction of cells (∼5-10%) which will be resistant to ferroptosis and/or less dependent on GPX4 for their survival. To our surprise, phenotypic characterization of RSL3 PDTPs in all TNBC cell lines (Basal, MSL and LAR) show a dramatic increase in cell-cell contact formation and closed colony growth patterns even in the MSL subtype that is not known to grow in colonies with cell-cell contacts (Fig. 6 A). As it can be well appreciated from figure 6A, this RSL3 PDTP phenotype is typically the opposite of chemotherapy driven PDTPs, where cells lost the cell-cell contacts in DTP development. We then investigated the expression of a panel of markers that are altered in proliferating chemotherapy-tolerant TNBC using IF staining and confocal microscopy analysis in PC, PDTP-P and PDTP-RSL3 of MDA-MB-468 cells (Fig. 6B) and quantitated the effect (Fig. 6C). As seen in the figures, PDTP-RSL3 of MDA-MB-468 cells show a robust buildup of E-cadherin at cell-cell contact junctions, with negligible detected vimentin and LC3B positive structures compared to PC. Moreover, PDTP-RSL3 cells show a mild increase in cytoplasmic expression of GPX4 (Fig. 6B, C). To complement the findings from IF analysis, we also performed WB analysis for these specific marker proteins (Fig. 6D) in MDA-MB-468 and HS578T cells and quantified the expression changes in folds of PCs (Fig. 6E). Western blotting results show an increase in GPX4 in PDTP-RSL3 in both the cells types. These results indicate that PDTP-RSL show more epithelial type morphology with increased GPX4 expression compared to PDTP-P and PC.

**Figure 6:**
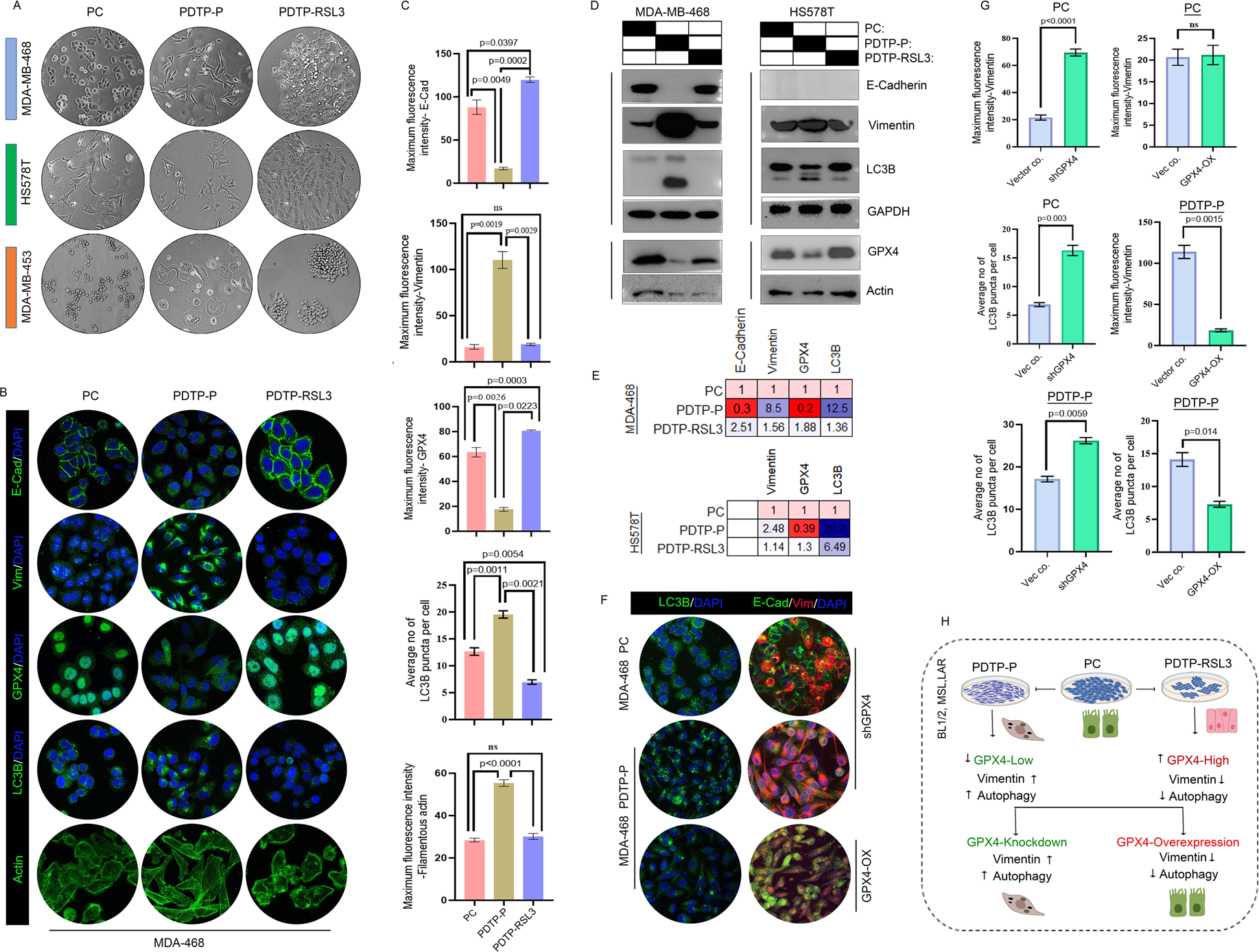
RSL3-PDTPs and modulation of GPX4 expression in TNBC PCs and PDTPs recognize GPX4 as a regulator of autophagy and EMT phenotype in TNBC. (A) TNBC cells from different subtypes were treated with RSL3 (1-3μM) for 72h to select RSL3 tolerant cells and were grown till recovery of proliferating colonies (15-20 days) of PDTP-RSL3 cells. Phase contrast images of morphological changes in PDTP-P and PDTP-RSL3 as compared to PC of indicated TNBC cell lines. Magnification: 10X. (B) Representative immunofluorescence staining of PDTP-P, PDTP-RSL3 and PC of MDA-MB-468 cell lines to assess the expression and localization of E- cadherin, vimentin, GPX4, LC3B and Actin (in green) and nuclear staining (Blue) with DAPI was performed. (C) Fluorescence intensity of different protein shown in B was quantified using FIJI ImageJ and presented as mean ±SD in the bar graphs with statistical significance between PC, PDTP-P and PDTP-RSL3. (D) The expression levels of E-cadherin, Vimentin, GPX4, and LC3B in TNBC cell lines and PDTP-P, PDTP-RSL3 and PC were analysed using western blots. (E) ImageJ software was used to quantify the band intensities of E-cadherin, Vimentin, GPX4, and LC3Band fold changes as compared to PC were presented as heatmap. (F) Representative confocal images of immunofluorescence staining of LC3B (Green), E-cadherin (Green) and Vimentin (Red) in PDTP-P and PC of MDA-MB-468 cell line in GPX4 Knockdown or GPX4 overexpression conditions are shown. (G) Fluorescence intensity of different protein stained in F were quantified using FIJI ImageJ. The expression of respective proteins as indicated in GPX4 knockdown and overexpression conditions in PDTP-P and PC MDA-MB-468 were compared with vector control. (H) Proposed model for GPX4 expression and its effect on EMT and autophagy summarised using the schematic.

In order to further verify if the phenotypic change that are observed in PDTP-RSL3 are strongly associated with an increase in GPX4, we performed shRNA media knockdown of GPX4 and over-expression of wtGPX4 in PC and PDTP-P in MDA-MB-468 cells by transient transfection methods. Immunofluorescence staining specific for LC3B, E-cadherin and vimentin expression and confocal microscopy analysis of GPX4 altered cells (Fig. 6F, G) indicate that transfection of GPX4 shRNA in PC resulted in a sharp increase in vimentin expression while GPX4 overexpression in PDTP-P resulted in a significant decrease in vimentin expression while increase in E-cadherin expression. Further, GPX4 shRNA transfection in both PC and PDTP-P resulted in the increase of LC3B positive puncta (Fig. 6F, G) and overexpression in PDTP-P decreased LC3B autophagy puncta significantly at the same time. Collectively, these results clearly suggest a strong coregulation of epithelial/mesenchymal behavior and autophagy flux by GPX4 protein in TNBC cells (Fig. 6H). Most importantly, these results highlight a strong ferroptosis/glutathione driven connection of cellular plasticity and autophagy in PC and PDTP TNBC cells that can be utilized as a signature to predict and/or detect drug tolerance in TNBC tumors under chemotherapy, apart from susceptibility to GPX4 and autophagy inhibition.

### GPX4-Vimentin shows an inverse correlation, and together with autophagy markers can predict survival in chemotherapy treated TNBC patients with high grade and metastatic tumors

Given the established association of GPX4 suppression, increased EMT and autophagy in TNBC PDTPs, we access the clinical relevance of our findings in Triple Negative Breast Cancer patients using different publicly available matched data. First, we asked if individual gene expressions of GPX4, vimentin (VIM) and autophagy marker LC3B2 (MAP1LC3B) have any implications on the recurrence-free survival (RFS) and distant metastasis-free survival (DMFS) in TNBC patient cohorts that corresponds to TNBC PDTPs. For this analysis, we have included the TNBC patient cohort form TCGA database that are treated with chemotherapy agents and have high grade disease with lymph-node metastasis (LN+). In this selected TNBC patient cohort we analyzed RFS and DMFS using KM-ploter for survival Kaplan Meier plots. As seen in figure 7A, a significantly decreased RFS as well as in DMFS is observed in patients with low GPX4 gene expression with a hazard ratio (HR) of 0.49 and 0.37, respectively. These results were also very consistent with GPX4 protein expression in TNBC as a marker for superior overall survival (OS), and DMFS in Liu_2014 proteomic dataset (Fig. 7B). We also analyzed and seen that GPX4 expression in general is a predictor of RFS in all the breast cancers treated with Neo-adjuvant chemotherapy (NACT) and DMFS in TNBC irrespective of grade and LN status with significant values (Fig. S4A), indicating that low GPX4 in associated with poor survival in patients that have aggressive and metastatic disease, analogous to the features enriched in TNBC PDTPs. Next, we analyzed the gene expression of VIM (Fig. 7C) and MAP1LC3B (Fig. 7D) for RFS and DMFS in TNBC cohorts with features of PDTPs, and interestingly, both the marker genes are strong predictors of poor survival individually with a significantly high HR value (>2) in LN+ high grade TNBC patients under chemotherapy. As indicated in Figure S4B, vimentin expression is also a predictor of poor RFS all breast cancer patients with NACT as well as for DMFS in TNBC with significantly high hazard values (Fig. S4B). Together these survival analyses suggest that VIM and GPX4 can strongly predict the survival of TNBC patients with chemotherapy and show reverse trends which are precisely aligned with our findings in cellular modeling of chemo-tolerance with TNBC PDTPs.

**Figure 7.**
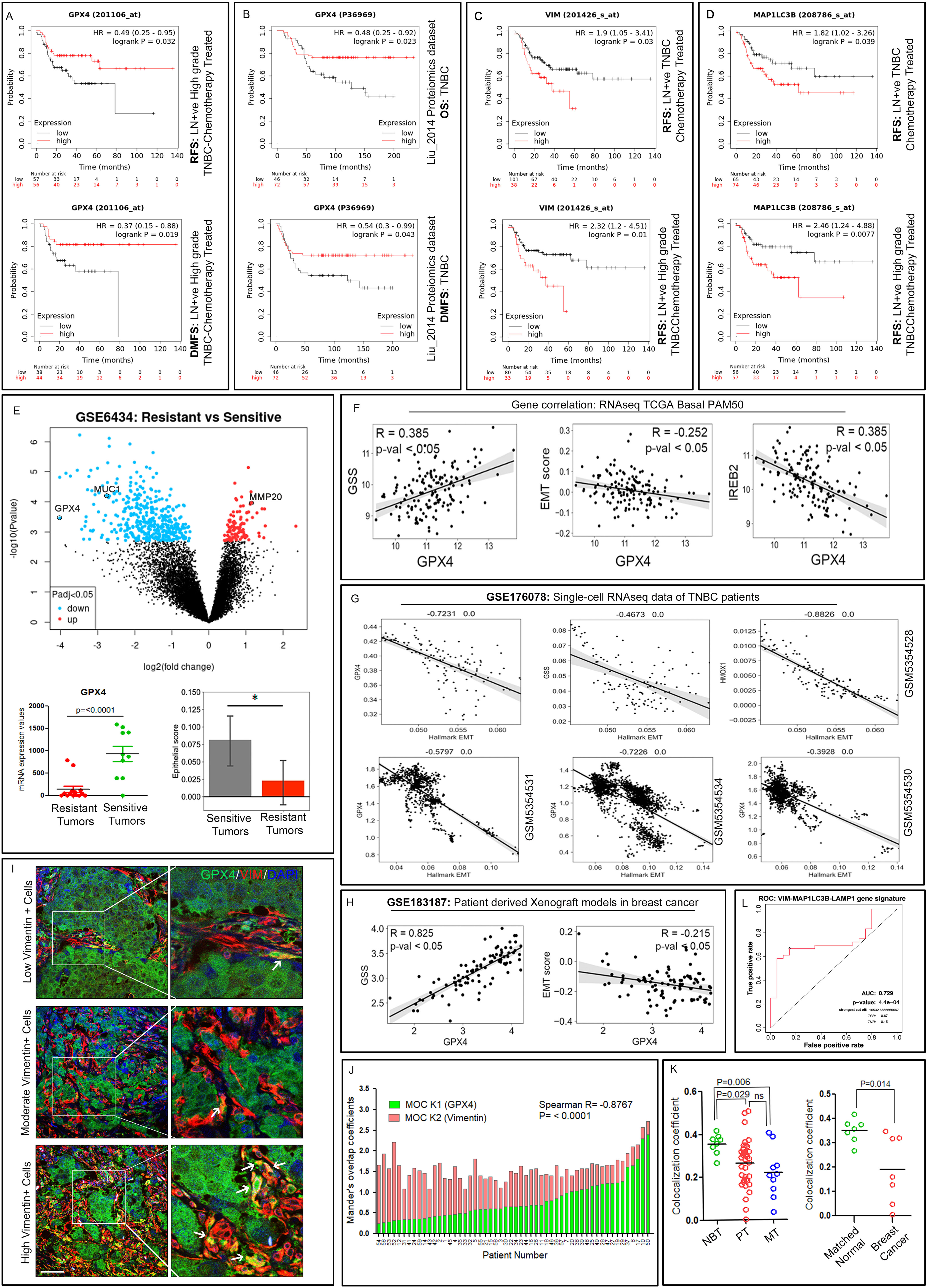
Low GPX4, high VIM and autophagy markers are strong indicator of worst outcome in chemotherapy treated TNBC patients with high grade and lymph node metastasis. Kaplan-Meier Plot representing relapse free survival and distant metastasis free survival in LN+ve high grade TNBC chemotherapy-treated patients with high or low gene expression (A) of GPX4 (C) vimentin (D) MAP1LC3B. (B) Kaplan -Meier Plot representing overall survival, relapse free survival and distant metastasis free survival in TNBC patients with high or low GPX4 protein expression from Liu_2014 proteomics dataset. (E). Volcano plot showing Differentially Expressed Genes analysed from gene expression Affymetrix dataset (GSE6434) using GEOR on GEO having breast cancer patient cohorts with docetaxel sensitive and resistant tumors. Significantly different and relevant genes were marked on the plot showing fold change and p value (Log2 FC and Log10 p value). Distribution of GPX4 gene expression between resistant and sensitive tumors is plotted in the dot plot below. Epithelial score was calculated as described under method section for the docetaxel resistant and sensitive tumors and plotted in the bar graph. (F) Correlation analysis showed GPX4 expression had a correlation with the GSS, EMT signature genes and IREB2 expression. (G) Scatter plot showing correlation analysis between GPX4 and EMT signature genes. Black line represents linear regression; grey area indicates the limits of the confidence intervals. (H) Correlation analysis of GPX4 with GSS and EMT score in patient derived xenograft models in breast cancer. (I) Confocal micrographs showing representative three channel overlap of immunofluorescence co-staining of GPX4 (green) and vimentin (red) proteins in paraffinized human breast cancer tissue sections, DAPI (blue) was used for nuclear staining. Areas marked under white squares were shown as zoomed inserts in the right panel where white arrows indicate cells showing co-expression of GPX4 and vimentin. Magnification, 63x and scale bar, 50 μM. (J) Waterfall bar graph showing the Mander’s overlap coefficients (MOC) for relative contribution of green (GPX4) and red (vimentin) channel signals to the colocalised immunofluorescence signals in 54 human breast tissues. Spearman’s R correlation between the groups was used to compute the correlation between MOC K1 and K2 and p value was determined by two-tailed t-test. (K) Dot blots (left) showing analysed colocalization coefficient for relative co-expression of GPX4 (Green channel) and vimentin (red channel) in human breast tissue used for immunofluorescence co-staining. Each dot in the plot represents an individual breast tissue categorised under normal breast tissue (NBT, n=8), primary tumor (PT, n=36) and metastasis tumor (MT, n=10). Dot blot (right) showing the colocalization coefficient for GPX4 and vimentin co-expression in tissue sections of human breast cancer (n=7) and its matched normal tissue (n=7) after immunofluorescence co-staining and analysis. Unpaired two-tailed t-test was used to analysis the significant difference between two groups and p values are indicated on the graphs. (L) ROC curve analysis of the 3-gene signature in LN+ high grade TNBC patients treated with chemotherapy as performed using ROCPlotter, https://www.rocplot.org/.

To further validate our experimental results, we analyzed a breast cancer gene expression Affymetrix dataset (GSE6434) on Gene Expression Omnibus (GEO) having breast cancer patient cohorts with docetaxel sensitive and resistant tumors using GEOR. Differentially expressed gene (DEG) analysis as seen in volcano plot in figure 7E drawn for resistant vs sensitive tumors shows GPX4 as the second most downregulated gene (-4.02 folds, Log2 FC; p value =3.47E-04, log10). Evaluation of key regulatory genes related to mesenchymal state (MUC1, TGFB1, MMP20, MMP3 and MMP16) (Fig. S4C) that are differentially expressed between resistant and sensitive tumors suggest that resistant tumors have more EMT-like state, that are very much aligned with our in-vitro findings (Fig. 2,3). Similarly, we found that expression of genes responsible for cellular ROS management (NFE2L2 and GSS) and iron binding, FTH1 were also compromised ROS in resistant tumors. Further, using the same dataset, we calculated the epithelial score (Fig. 7E) and found it to be significantly lower in resistant tumors compared to tumors in the sensitive cohort. Collectively, this analysis supports the idea that chemo-resistant tumors have most likely gained a mesenchymal state (Fig. 2) and have compromised ROS handling capacity (Fig. 5).

In order to map the clinical correlation of GPX4 with markers of ferroptosis, epithelial and EMT states, and iron homeostasis, we have analyzed several bulk tumor gene expression databases (TCGA, METABARIC, Wang, Lu and Ivshina) for different breast cancer types, Basal-Like bulk RNAseq data from TCGA, single-cell RNA sequencing datasets from TNBC patients as well as gene expression sets from TNBC PDX models as shown in figures 7F-H and S4D. Gene expression correlation dot-plots in different breast cancer datasets show that GPX4 expression is largely positively correlated with epithelial type cytokeratin genes (KRT8, KRT18 and KRT19), while negatively correlated with mesenchymal, metastasis markers and EMT regulator genes such as VIM, S100A4, CDH2, MMPs, SNAI2, ZAB2 and MET (Fig. S4E). Further, we have observed that GPX4 shows significant positive correlations with GSS gene, while significant negative correlations were seen with EMT score as well as IREB2 in TCGA RNAseq Basal-Like PAM50 cohort (Fig. 7F). Single-cell RNAseq data from individual TNBC patients (GSE176078) also highlighted robust negative correlations of GPX4 with hallmarks of EMT (Fig. 7G) in all the patients, indicating a very high clinical relevance of our finding in PDTPs that GPX4 can differentially regulate epithelial and mesenchymal cellular behaviors (Fig. 6). Very similar trends for GPX4 were also re-capitulated in the patient derived xenograft models in breast cancer from GSE183187 (Fig. 7H). Altogether, these data show a robust association of GPX4 with EMT phenotype across TNBC cell lines, human tumor samples and PDX models which has not been mapped earlier in breast cancer.

Since we have seen a strong inverse relation of GPX4 and VIM at gene expression level, we sought to investigate this correlation on protein expression levels in breast cancer to establish a functional connection between these proteins in normal breast, primary and metastatic breast tumor tissues. In order to analyze expression of GPX4 and vimentin together in human tissues, we performed dual immunofluorescence co-staining of paraffinized breast tissue sections in each type (Fig.7I). We have analyzed in total 54 breast tissue sections having normal breast tissue (NBT, n=8), primary breast tumors (PT, n= 36) and lymph node metastases (MT, n=10). As shown in the confocal micrographs of immunofluorescence co-staining of GPX4 (green) and vimentin (red) in representative breast cancer sections tumor sections (Fig. 7I), a mutually exclusive staining of these two proteins in tumor cells were observed. Tumor sections with different degrees of vimentin positive cells in-between well differentiated epithelial tumor mass largely positive for GPX4 was observed, with few cells strongly co-expressing both the proteins. In order to substantiate these findings on GPX4-vimentin staining patterns we analyzed the co-localization coefficient to detect extent of their co-expression in cells of breast tissue sections (Fig. 7K) and their relative contribution in the co-localized immunofluorescence signals by estimating the Mander’s overlap coefficient (MOC) (Fig. 7J). As evident from the overlay images of the red and green channels of immunofluorescence co-staining in the breast tissue sections (Fig. 7I), analysis of MOC K1 and K2 showed a very strong and significant opposite correlation spearman’s correlation, -0.8767, with p<0.0001 (Fig. 7J) suggesting that protein expressions of GPX4 and vimentin share a negative correlation and are minimally expressed. These results are intensely in line of our cellular and in-silico results. Further, we also analyzed the relative co-localization coefficients in normal and tumor tissues to understand if co-expression of these two proteins change with tumorigenesis and metastasis, as seen in the dot-blots in figure 7K, it is evident that normal breast tissues has significantly higher co-expression of GPX4 and vimentin than both, primary tumors and metastatic tumor tissues, while there is not much difference between the co-localization coefficients in primary and metastatic tumors, indicating that the changes in expression ratio between these protein can be predictive of breast tumorigenesis. These results were also strongly affirmed when we analyzed the co-expression of GPX4 and vimentin in matched tumor and normal breast tissue section (Fig. 7K). Finally, we also analyzed the mean florescent intensity of GPX4 and vimentin staining in all 54 breast tissue sections and found that GPX4 expression is significantly higher in these tissues as compared to vimentin (Fig. 4SE), while there is a higher expression of GPX4 in primary and metastatic tumors compared to normal breast tissue (Fig. 4SE). Together these findings suggest a negative correlation of GPX4-vimentin in breast tissue at protein expression which is suggestive of opposing roles and regulatory functions at cellular level as suggested by our experimental data (Fig. 6), apart from a identification of significantly increased co-expression in normal breast tissue, which might be important for the homeostasis of the normal breast tissue.

Further, as our preceding data analysis on human breast cancer establish a reverse equation between GPX4 and vimentin expression, and their expression can predict (RFS and DMFS) survival, especially in aggressive TNBC patients with chemotherapy treatment, we then queried if a gene signature comprised of mesenchymal marker, VIM- and autophagy markers MAP1LC3B and LAMP1 can be a better predictive biomarker for pCR and RFS in TNBC patients with chemotherapy. For that, we have performed the ROC (receiver operating characteristic) analysis with ROC plotter to compute pathological complete response and RFS in chemotherapy treated TNBC patients built on AUC (Area Under the Curve) values (*68*). As seen in figure 7L, expression of VIM-MAP1LC3-LAMP1 tri-gene signature can effectively predict 5-year RFS in TNBC patient with a statistically significant (p=4.44-04) AUC value of 0.728. These genes are highly co-expressed in chemotherapy non-responder TNBC patients (Fig. S4F) compared to responders. It was also seen that MAP1LC3B expression also could individually predict RF-survival in TNBC (Fig. S4H) but not to a decisive extent than the tri-gene signature (Fig. 7L). Likewise, we have also analyzed the ROC plot for GPX4-CDH1 gene signature which was found to be significantly (p=1.6e-02) downregulated in aggressive TNBC patients who do not achieve pCR with chemotherapy (Fig. S4G) with AUC value of 0.609. All these results from breast cancer patient samples, TNBC patient data analysis strongly suggest that the gene and protein expression alterations which were linked to development of TNBC PDTPs were also stood significantly valid in TNBC patients and are very strong predictor of chemotherapy response in patients who have metastatic and high-grade tumors, somewhat analogues to phenotypes of PDTPs that we have raised in-vitro studies.

## DISCUSSION

In the present study, we show that a minor portion of TNBC cells can enters into a latent cytostatic DTP state upon treatment with high doses of chemotherapeutic agents and eventually regain a proliferative PDTP state in the absence of drug. We found that though TNBC cells from all the four distinct subtypes can undergo a DTP state and attain proliferation with different classes of chemotherapeutic agents. Basal-Like subtypes entered into a fully non-reversible EMT, while the LAR subtype of cells could only achieve a partial EMT state with an increase in vimentin but retention of cell-cell contacts (Fig. 8A). Nevertheless, despite the phenotypic differences, all the PDTPs display acquired chemo-resistance with elevated levels of autophagy flux, glutathione depletion and lipid peroxidation that turned out to be vulnerabilities of the PDTP state in TNBC (Fig. 8B). Further, we show that the mesenchymal state and autophagy in PDTP cells are dependent on GPX4 expression status, as RSL3-tolerant cells with elevated GPX4 or ectopic overexpression of GPX4 in these cells can enhance junctional E-cadherin and suppress vimentin expression as well as LC3B positive puncta, while shRNA mediated depletion of GPX4 can induce vimentin expression in PC. Additionally, analysis of different TNBC patient databases suggests that GPX4 expression is positively correlated with epithelial markers while a negative correlation with EMT hallmarks. Most importantly, we establish an inverse correlation of GPX4 and vimentin expression in breast cancer. At both, gene and protein levels, GPX4 expression is significantly associated with better patient survival, especially in chemotherapy treated, high grade metastatic TNBC patients. Finally, we defined a signature of mesenchymal and autophagy genes, VIM-MAP1LC3B-LAMP1 that effectively predicts poor survival in these settings (Fig. 8C). Collectively, our findings reveal distinct and convergent cellular landscapes of the PDTP state in different TNBC subtypes that underscore the therapeutic opportunities to target shared vulnerabilities of aggressive TNBC cells with acquired proliferative chemotherapy-tolerant state.

**Figure 8.**
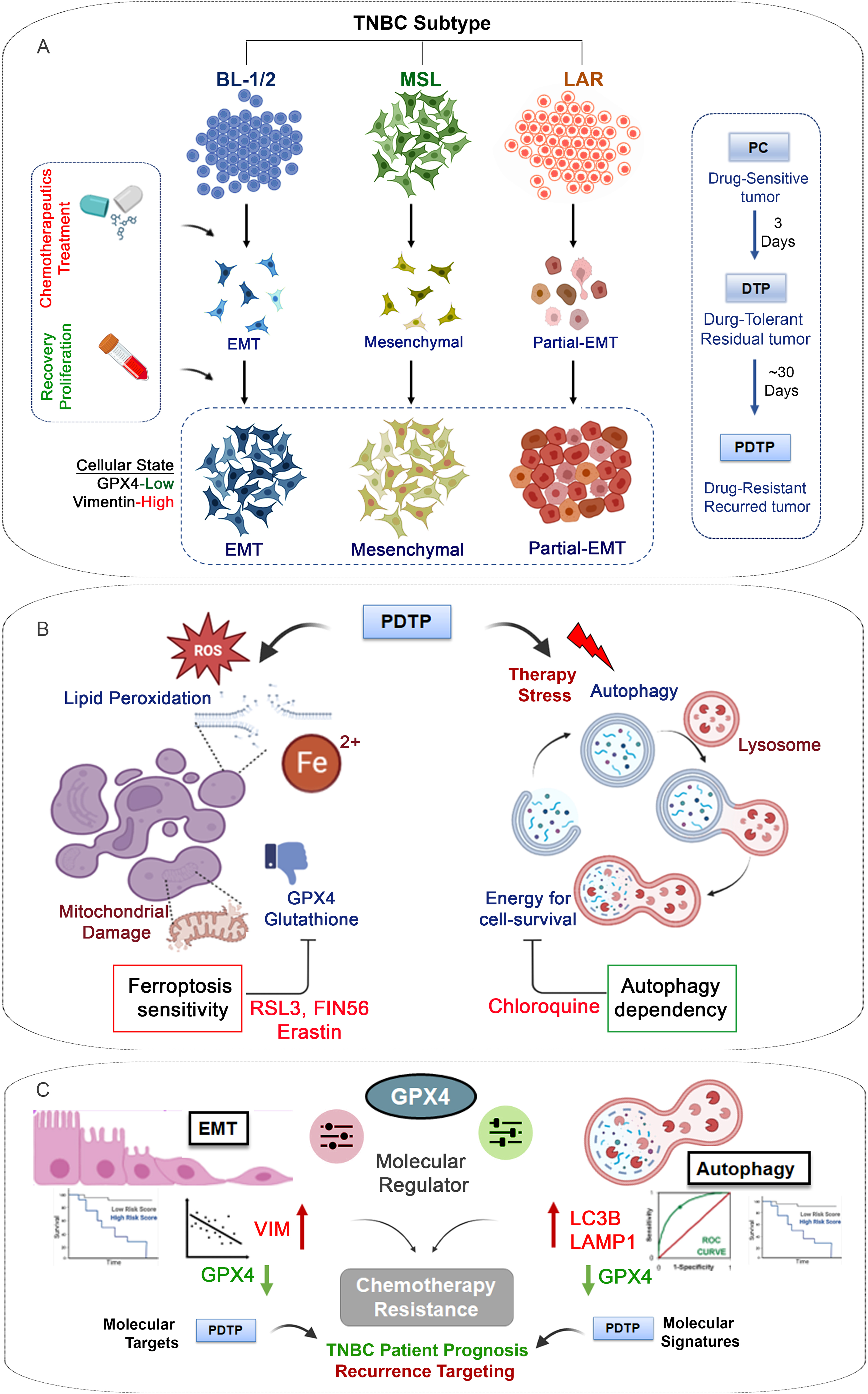
Schematic model summarizing the main findings of the study. **(A)** Derivation of PDTP from different subtypes of TNBC. **(B)** Identification of common targetable vulnerabilities of TNBC PDTPs **(C)** Regulation of PDTP phenotypes by GPX4 and its prognostic and therapeutic potential in TNBC patients.

Growing reports in various solid tumor types have substantiated that a sub-population of tumor cells that escapes targeted or chemo-therapeutic drugs can persist in a dormant state under therapy, and can regain proliferation that is clinically presented as post-therapy tumor recurrence with metastasis (*17*, *21*, *23*, *69–72*). These therapy-tolerant cells can evolve either due to the selection of pre-existing resistant clones or via a significant molecular reprogramming during treatment that results in a cellular adaptive response (*26*, *41*, *73*). Several studies have independently affirmed that TNBC tumors recurrence rates are substantially higher in patients with residual disease and residual tumor burden is a very strong indicator of worst event free survival in TNBC patients worldwide (*14*, *74–80*). Our cellular modeling of DTPs using different classes of chemotherapeutics supports the consolidated clinical observation as we have observed that TNBC cells have a high potential to overcome the drug induced stress condition as DTP and attain aggressive PDTP state, a feature that is independent of the molecular subtype. Moreover, it was seen that post-therapy recurrence in TNBC is mostly associated with more resistant and metastatic disease that accounts for worst clinical outcomes in these patients was also recapitulated in our PDTP modeling. We show that PDTP cells were more resistant to the initial therapy agent and show enhanced capacities of colony formation, migration and metastasis (*21*, *81–84*).

It has been widely accepted in clinical studies that drug resistance to chemotherapy mostly arises in a non-genetic manner in TNBC. Earlier studies investigating chemoresistance in TNBC patients utilizing a vanity of methods like *in situ* hybridization (*85*), profiling of bulk genomic (*86*, *87*) and tumor cytogenetic markers highlighted that neo-adjuvant therapy did not alter the tumor genetic diversity however, have enriched tumor cells with more mesenchymal phenotypes (*85*). These clinical research observations are in line with our findings that TNBC attains chemoresistance by acquiring a transient non-genetically driven DTP state. Further, PDTP cells indeed are more in a mesenchymal state with increased vimentin expression in all subtypes. Other recent studies focused on longitudinal tumor cell sampling in TNBC patients under neoadjuvant chemotherapy in addition to anti-VEGF antibody Bevacizumab (*88*) to analyses drug resistant residual tumors, suggest that non-mutational transcriptome changes are primarily responsible for the development of resistant cancer cells. Similarly, analysis of pre- and post-neoadjuvant chemotherapy treated tumor samples (*89*) as well as PDX derived chemo-resistant tumors (*90*) show that resistant genotypes and persister clonal populations with distinct transcriptome and proteome were pre-existing and may have enriched under therapeutic selection. All these previous studies and our current study suggest an evolution of the DTP population from pre-existing cell population with extensive molecular and phenotypic switching, including EMT adaptations. More interestingly, we have seen that Basal-Like TNBC subtypes can enter into a stable EMT state, however, the luminal cell enters a partial EMT state. It has been shown using genetic mice models that breast cancer cells that enter partial EMT have higher capability to initiate metastatic tumors, while complete EMT promote drug resistance (*44*). As tumors from LAR subtype are the most difficult to treat with chemotherapy and show higher metastatic rates, a more detailed understanding of this unique capacity to undergo transient EMT state in LAR tumors can elucidate the underlying mechanism of worst seen clinical response in these tumors (*45*, *91*).

Transcriptional reprogramming in DTPs can also offer altered metabolic states that are also among the mechanisms ensuring “persister cell” proliferation, invasion, and resistance to therapies. Lately, it has been shown that cells under stress and EMT has altered energy acquisition mechanisms, such as dependency on OXPHOS (*90*) and autophagy (*23*), for their survival and proliferation. Detailed sub-cellular characterization of TNBC PDTPs in our study highlighted increased autophagy in these cells required for their survival and such dependencies also pose therapeutic opportunities in these hard-to-treat residual disease. Likewise, cellular changes in reactive oxygen handling, iron and lipid metabolism are previously known in the DTP state for several solid tumors and have been implicated in the acquisition of cell resilience to iron-induced oxidative ferroptotic cell death (*67*, *92*). Such mechanisms allow cancer cells to resist apoptosis induced by chemotherapeutic agents (*18*). While it has been known that TNBC cells are vulnerable to iron chelators(*67*, *93*), ferroptosis inducers (*94*), and deprivation of anti-oxidant pathway metabolites (*33*), surprisingly, we observed that PDTP cells further enriched these deleterious metabolic aberrations with much higher states of lipid ROS, labile iron and low glutathione status. We found that TNBC PDTP cells have unvaryingly downregulated metabolic players like GPX4, and others like HO-1, xCT, and NRF2 at both gene and protein levels. Previous studies suggest that metabolic reprogramming might be required to acquire and maintain the DTP state and consequently higher cellular ROS that is also involved in intracellular signals for autophagy, proliferation, cytoskeletal remodeling, cell migration and invasion phenotypes, and in some cases also in extracellular matrix remodeling (*67*, *95*). These redox driven advantages on the other hand disposes TNBC PDTPs to a potentially lethal vulnerability affair if targeted using GPX4 inhibitors or ferroptosis inducer like xCT inhibitor (*66*, *96*).

GPX4 suppression has been reported in many solid tumors under DTP states as one of the essential molecular changes, however, if the DTP or PDTP states are phenotypically dependent on compromised glutathione state has never been clearly answered experimentally. Our study identifies that GPX4 is a robust regulator of autophagy and EMT in TNBC cells; pathways that are frequently reported to be upregulated collectively during metastasis in several cancer types (*57*, *60*, *91*). In light of these results, we speculate a potential anti-tumor role of GPX4 in TNBC cells, apart from regulating ferroptosis, especially in suppressing phenotypes such as metastasis and drug resistance. Interestingly, our in-silico data analysis using several TNBC patients’ datasets strongly advocate association of low GPX4 with worst survival in TNBC patients as well as a significantly negative correlation of GPX4 with EMT and metastatic markers like VIM. Even though, it is usually challenging to estimate autophagy flux in tumors using gene expression analysis, strong stand-in markers, such as MAP1LC3B and LAMP1/2 are shown to be unswervingly correlated with increased levels of autophagy in diverse tumor types (*97–99*) and in some of them with poor survival and metastasis (*100–103*). More interesting, we have exploited a tri-gene signature, VIM-MAP1LC3B-LAMP1 that combines EMT, autophagy, and lysosomal abundance phenotypes and could efficaciously predict response to chemotherapy in patients with high grade TNBC tumors with metastasis.

In conclusion, the present work primarily expands the understanding on distinct and shared phenotypes and vulnerabilities of proliferating DTPs from different TNBC molecular subtypes that are representative cellular models of recurrence associated drug-resistant residual subpopulation in patients. There is a very limited understanding on how dormant and proliferation states in TNBC DTP develop from chemotherapeutic agents in general (*23*). Despite the fact that TNBC are not only highly heterogenous but also shows the highest incidences of recurrence and chemoresistance among all the types of breast malignancies, comprehensive and comparative studies based on modeling of TNBC subtypes and drug specific DTPs have not been established. Our study has acquired systematic experimental understanding in this direction and laid a platform to address several significant questions on TNBC DTPs and their intricate behavioral adaptations to survive and overcome severe chemotherapy-induced stress conditions that would equip us to design precise molecular approaches to target this PDTP state. A substantial degree of in-depth studies on TNBC DTP/PDTP states are further required in order to clinically translate the tangible strategies against the DTP state of cancer cells in patients. These efforts will be more in lines of establishing persuasive patient derived organotypic and mice models with representation of TNBC subtypes with molecular profiling and their extensive exploitation using mathematic modeling-based tools to discover unique intrinsic/extrinsic dependencies that can transcribe synthetic lethal therapeutic combinations to eliminate the DTP and PDTP cells.

## Materials and Methods

### Chemicals, Drugs and DNA constructs

Chemotherapeutic agents used in the study were purchased from Sigma (Paclitaxel, catalog no. T1912; doxorubicin, catalog no. D1515 and cisplatin catalog no. PHR1624, GPX4 inhibitors RSL3, catalog No. SML2234 and Fin 56 catalog no- SML1740). xCT inhibitor Erastin catalog no: E7781. Thiazolyl Blue Tetrazolium Bromide (MTT) (M5655), DAPI (4’,6-diamidino-2-phenylindole) and Chloroquine (catalog no. C6628) were procured from Sigma-Aldrich. Matrigel was purchased from Corning. GPX4 shRNA in cloned in PLKO.1-puro-shRNA plasmid were purchased from Sigma-(TRCN0000046249, sequence: 5’GTGAGGCAAGACCGAAGTAAA3’ and TRCN0000046251, sequence: 5’GTGGATGAAGATCCAACCCAA3’). GPX4 overexpression vector- 442-PL1 wad provided as a kind gift by Prof. Dr. Bettina Kempkes (Ludwig –Maximilian-University, Munich, Germany

### Antibodies

The following antibodies were purchased from Cell Signaling Technology (USA) p62 catalog no. 5114T, LC3B catalog no: 2775S, E-cadherin (610812) from BD bioscience and mouse Vimentin from sigma catalog no: V6389, following antibodies were purchased from abcam HO-1 catalog no: ab137749, SOD1 ab52950, GPX4 catalog no. ab125066. HRP-conjugated IgG anti-rabbit (catalog no. A21234) and anti-mouse (catalog no. A28177) were purchased from Life technologies. Alexa fluor 488 (anti-rabbit catalogue no. A11008 and anti-mouse catalogue no. A11001) and 568 (anti-mouse A11004, and anti-rabbit A11011) were purchased from Invitrogen.

### Cell culture

Different subtypes of triple negative breast cancer (TNBC) cell lines MDA-MB-468, MDA-MB-231, MDA-MB-453, HCC70, and HS578T were originally procured from the American Type Culture Collection (USA). These cells were maintained in RPMI medium (Gibco BRL, USA) in which l-Glutamine (2 mM) and sodium bicarbonate (2g/liter) were added. The media was supplemented with 10% fetal bovine serum (Gibco BRL, USA) and penicillin/ streptomycin (Invitrogen, USA). The Cells were cultured in 5% CO_2_ humidified incubator at 37°C. All the cultures were routinely checked for mycoplasma using a commercially available kit form Biological Industries, Israel and DAPI nuclear staining.

### Generation of DTP and PDTP TNBC models

Cells were treated with increasing doses of chemotherapeutic drugs (Paclitaxel, Doxorubicin and Cisplatin) for 72 h and cell viability assays were conducted, IC_50_ values were determined for each drug using Graphpad 5.0 software as described previously (*35*). To generate DTPs IC_85-90_ concentrations of each drug was used to treat the cells for 72 h and thereafter let the residual drug-tolerant persister cells recover till they growth by maintaining them in drug free medium.

### MTT assay

To determine the cytotoxic effect of drugs, dose response experiments were performed. In a 96 well plate format MDA-MB-468 (12000 cells), HCC70 (9000 cells), MDA-MB-231 (9000 cells), HS578T (3000 cells) or MDA-MB-453 (9000 cells) cells were seeded for 16 hours. Cells were treated with drugs for 72 hrs with an increasing concentration of drugs. Cell viability were determined using MTT assays as described previously (*35*), till the visible formation of blue formazan crystals. After solubilized of formazan crystals using 100 µl of stop solution, the Optical Densities (OD) were determined at 570 nm (reference wavelength 630 nm) using microplate reader Cytation5 spectrophotometer (Biotek, Agilent). Dose response curves were generated using Graph-Pad Prism 5.0 software and IC_50_ were determined.

### Colony formation assay

Cells were seeded up to 5, 10 and 15% confluence and allowed to grow to form visible colonies. Media was replenished after every three days. Colony was fixed in 4% PFA and stained with crystal violate. Photographs of the plates were taken a 16.2 Megapixel digital camera (Nikon D7000) using light box to illuminate the plate.

### Transmission Electron Microscopy (TEM)

To study the sub-cellular details using TEM, cells were grown as a confluent monolayer, fixed with 3% glutaraldehyde, washed with 0.1 M of sodium cacodylate pH 7.4, and postfixed with 1% osmium tetraoxide. Cells were then dehydrated and processed for sample preparation. Grids were contrasted with alcoholic uranyl acetate for 30 seconds and dipped once in lead citrate. Under a JEOL 1400Plus transmission electron microscope, the grids were observed at an accelerating voltage of 120 KeV and 10000X. Images were then captured using a Slow Scan CCD camera (TRS, Germany).

### Immunoblotting

Whole cell lysate of different cell lines for total protein was prepared in lysis buffer consisting 0.2% Triton X-100, 50 mM HEPES (pH 7.5), 100 mM NaCl, 1 mM MgCl2, 50 mM NaF, 0.5 mM NaVO3, 20 mM β-glycerophosphate, 1 mM phenylmethyl sulfonyl fluoride, leupeptin (10 μg/ml), and aprotinin (10 μg/ml) as described previously (*35*). Total protein concentration of the cell lysate was estimated by Bradford assay (Bio-Rad, Hercules, CA). For Western blotting equal amounts (up to 80 μg) of total cell lysates were loaded on SDS–polyacrylamide gel electrophoresis system (Bio-Rad) and transferred on nitrocellulose membrane which was proceeded for immunoblotting to detect specific proteins. The membrane was probed with Primary antibodies for GPX4, xCT, HO1, SOD1, p62, LC3B, E Cadherin, Vimentin (dilution, 1:1000), and GAPDH/alpha-tubulin/beta actin (1:2000) as a loading control. After incubation with species specific secondary HRP-labelled IgG (1:5000), membranes were developed using ECL reagent (Bio-rad) and the images were acquired on ChemiDoc Imaging System (Bio-Rad). Images were used for the densitometric analysis using the ImageJ software (National Institutes of Health, USA), and the intensity of each band was normalized to its respective loading control. The data was presented as folds of control.

### Isolation of total RNA and real-time PCR reactions

Cell lysates were prepared in RNA later buffer and Qiagen RNeasy Kit were used to isolate total RNA as per the manufacturer’s instructions. The cDNAs were transcribed from 2µg of RNA using ABI High-Capacity Reverse Transcriptase Kit (Applied Biosystems, Life Technologies). Ten nanograms of cDNA were used for the real time PCR using ABI SyBr Green PCR Master mix (Applied Biosystems, Life Technologies) in QuantStudio 12K Flex Real-Time PCR System (Life Technologies) with gene specific primers sets, β actin or GAPDH gene expression levels were used as for normalization in all experiments. The expression of target genes was plotted as fold change of experimental controls after calculating the relative to GAPDH or Actin as a housekeeping gene using the 2(-ΔΔCt) method. The sequences of forward and reverse oligonucleotides used in this study are shown in Table S1.

### Migration assays

Cells were seeded in 96 well plates at 90-100% confluence and serum starved overnight. cells were treated with 5µg/ml of Mitomycin C for 3 hrs. The scratch was made using a 200µl sterile tip and cells were stimulated with 10% FBS containing complete medium. Phase-contrast images of migrating cells at initial and different time points were captured using inverted microscope (Optika-SRL, IM-5FLD, C-P6 fitted with 6.3 Megapixel camara) at 20x objective. The average distanced travelled were quantitated from 3 independent experiments. Data was plotted in percent wound closure and migration rate with time.

### Invasion Assay

Matrigel degradation and invasion assays were performed using Boyden chambers (BD Falcon) of 8 µm pore size in a 24 well format. Boyden transwell chambers were coated with 5µg of the Matrigel suspended in 100µl of the ice-cold media and incubated at 37°C for 1 hr. In 200µl of serum free media, 2 x10 ^5^ cells were seeded in the Boyden chambers and in the lower chamber 450µl of media containing 10% of FBS were added to incubate for 16 hrs. The Boyden chambers were taken out, washed in PBS and fixed in 4% Para-formaldehyde (PFA) in PBS for 10 min. The chambers were than washed in PBS and stained in 0.03% crystal violet solution for 10 min. Cells from the inner surface of the chamber were carefully removed using cotton ear buds and chambers were washes with PBS and air dried. Images of the lower surface containing invaded cells were acquired from 10 different fields on an upright AxioImager Z1 microscope (Carl Zeiss, Jena, Germany) using a 10x objective lens. Number of cells invaded were calculated from 3 independent experiments and data was plotted as folds of control.

### Immunofluorescence staining

Immunofluorescence staining of cells was performed as described earlier (*104*). Briefly, cells grown on coverslips and treated as per the experimental procedures, the cells then washed with PBS and either were fixed using 4% PFA in PBS (20 min at RT) or in chilled methanol (10min at -20° C). After the fixation, the cells were washed 3 times with PBS and permeabilized with 0.25% triton x 100 in PBS for 10 mins. Cells were incubated with primary antibody at 4°C for overnight followed by PBS washes 3 times. Alexa 488 and 560 Secondary antibody incubation was performed for 1 hr at room temperature. Nuclear staining was done with 4′,6-diamidino-2-phenylindole (DAPI, Sigma). Confocal microscopic images were acquired using Leica confocal microscope at 63x oil emulsion objective using Leica Application Suit X (LAS X) software platform. Images were processed and analyzed using LAS X software.

### ROS measurements

To quantitatively measure intracellular reactive oxygen species in live cell samples we have used cell permeant reagent 2’,7’–dichlorofluorescin diacetate (H_2_DCFDA) from Cellular ROS Assay Kit (Abcam, ab113851) as per the manufacturer’s instructions. For the experiment, cells were cultured in a black walled clear bottom 96-well plate for overnight. The cells then were treated with ferroptosis inducer as indicated in figures. For estimation of ROS, Cells were then stained with 10 μM H_2_DCFDA for 30 min at 37°C protected from light. After washing the cells, the plate was read on a microplate reader (Cytation5 spectrophotometer, Biotek, Agilent) upon setting fluorescence at Ex/Em = 485/535 nm. Data are presented as folds of controls. The reading values were normalized to the number of living cells in each well.

### GSH measurements

Total GSH levels in cells were detected by using luminescence-based GSHGlo Assay reagent kit (Promega, catalogue no. B6912) as instructed by the manufacturer. The assay kit detects and quantifies total glutathione (reduced and oxidized forms) in the cell lysates. In order to detect GSH cell lysates of control and treatment experimental sets were prepared directly in the multiwall plate by adding 50µl GSH specific lysis buffer with the reagents provided in the kit followed by a 5 min incubation and then added 50µl of Luciferin Generation Reagent in each well. The mixture was incubated for 30 mins at RT and assays are mixed and incubated for 30 minutes. After the incubation, 100µl of Luciferin Detection Reagent was added to all wells and mixed properly. After a 15-minute incubation, luminescence is measured using a multiplate reader (Cytation5 spectrophotometer, Biotek). GSH content was calculated directly from luminescence measurements and the data was presented as means ± SD of fold of controls.

### Iron measurements

For measuring cellular iron in Fe^2+^ and Fe^3+^ states as well as total iron content we have a used a colorimetric Iron Assay Kit system (Abcam, ab83366) in stepwise manner as described in the manufacture’s protocol. Briefly, up to 2.5 x 105cells were grown and treated in 6 well plates, post treatment cells were washed with cold PBS and then lysed in 100 μl of iron assay buffer. Cell lysates (100 μl) were incubated with 5 μl of iron reducer for 30 mins at 37°C in a 96-well plate format. After the incubation, 100 μl of iron probe was added to detect ferrous and ferric ions. The absorbance was immediately measured at 593 nm with a microplate reader (Cytation5 spectrophotometer, Biotek). Results were normalized to protein content in cell lysates and expressed as means of folds of control.

### Lipid Peroxidation assay

A florescence shift-based assay (Image-iT Lipid Peroxidation Kit, Cat. No. C10445, Thermo Fisher Scientific) was used to analyze the peroxidation of lipid in live cells exploiting the BODIPY 581⁄591 C11 fluorescent reporter lipid probe as described previously (*35*). Cells were plated in black wall clear bottom 96-well Microplates (Corning, cat. No. 3603) for 16 hrs, proceeded for experimental control and treatments as indicated. The cells were then incubated with 10 μM lipid peroxidation sensor C11-BODIPY (581/591) in complete medium and incubated for 30 min in a humified incubator with 5% CO2 at 37°C temperature. Cells were washed with PBS and 100μl of fresh medium was added on the cells. The cells were then incubated in the live cell imaging system (IncuCyte S3, Sartorius) for acquiring bright field and florescence imaging at 10x magnification using a set of excitation/emission of 581/590 nm (Texas Red filter set) for the reduced dye, and the other at an excitation/emission of 488/510 nm (fluorescein isothiocyanate filter set) for the oxidized dye. The red-to-green fluorescence shift was used to detect lipid peroxidation in acquired images and the data was analyzed using IncuCyte 2020A software.

### Tissue Array Immunofluorescence

Human breast cancer tissue array (catalogue number: NBP2-30212, NOVUS) were deparaffinized and hydrated as per the manufactures’ instruction which was followed by antigen retrieval in heated tris-EDTA buffer (pH-9) using microwave, and then CuSO_4_.NH_4_Cl (pH-6) mediated auto-fluorescence quenching for 15 minutes. Tissue array were permeabilized using 0.2% triton X-100 in tris buffer for 15 minutes and stained with primary antibody for vimentin and GPX4 in 1:100 dilution. Fluorophore tagged secondary antibody (Alexa Fluor 568 anti-mouse IgG of Invitogen, A-11004 and Alexa Fluor 488 anti-rabbit IgG of Invitrogen, A-11001) were used in 1:200 dilution and incubated for 1 hrs at room temperature. DAPI was used for nuclear staining and coverslip was mounted using Mowiol based mounting medium. Immunofluorescence images were acquired on 60x objective of Nikon’s 10^th^ generation AX/AX R confocal microscope and quantitation of mean fluorescence intensities in each channel and colocalization coefficient (Pearson’s colocalization coefficient) and contribution of each channel to the colocalization was estimated using Mander’s overlap coefficients k1 and K2. These parameters were used for scoring the tissues staining using Nikon elements AR analysis 5.42.00 software to estimate colocalizing coefficient. Data was presented in mean florescence intensity from three representative images per tissue section in each channel and their contribution to the colocalization.

### Bioinformatic analysis

#### Pre-processing of Datasets

The Microarray (GSE6434) dataset was retrieved from NCBI GEO database and was pre-processed to procure the gene-wise expression from the probe-wise expression matrix using respective annotation files for the mapping of probes to genes. For single-cell RNA sequencing data (GSE176078), the MAGIC (version 2.0.3) (*105*) imputation algorithm was used to recover noisy and sparse single-cell data using diffusion geometry. Appropriate platform annotation files were utilized to map individual reads to genes. RNAseq (GSE183187) raw counts were obtained from NCBI GEO and was used to align the reads with hg38-human (or mm10-mouse) reference genome. They were then normalized for gene length and transformed to TPM (transcripts per million) values. Log2 normalization was carried out to acquire the final expression data. TCGA datasets were obtained using UCSC Xena tools.

#### Scoring Methods

To quantify the enrichment of epithelial and mesenchymal signatures, ssGSEA (single sample gene set enrichment analysis) was carried out on KS epithelial and KS mesenchymal gene lists, respectively, using the GSEAPY python library. The normalized enrichment score (NES) for these gene sets was obtained wherein a higher Epi score denotes a more epithelial phenotype, whereas a higher Mes score indicates the enrichment of a mesenchymal phenotype. The EMT score was calculated by subtracting the Mes score from the Epi score. The gene set for Hallmark ROS pathway, Hallmark Apoptosis pathway and Hallmark OXPHOS pathway was obtained from MSigDB (*106*). These gene sets were used to calculate the respective scores for the pathways using ssGSEA method. The activity scores of Hallmark pathways and EMT signatures for the single-cell RNA sequencing datasets were computed using AUCell (version 1.18.1) (Aibar et al. 2017) (Aibar et al. 2017) from the R package ‘Bioconductor’ with default parameters.

### Datasets

Gene and protein expression analysis of breast cancer patients were done based on The Cancer Genome Atlas (TCGA) breast cancer dataset. For Survival analysis TCGA gene expression was used on Kaplan-Meier Plotter (https://kmplot.com/analysis) and Recurrence Free Survival (RFS) was analysis using median cut. For protein expression TCPA portal (https://tcpaportal.org/tcpa/survival_analysis.html) was used to derive survival, and Kaplan-Meier plots were derived with BRCA data set containing 901 patient samples.

ROC plots in chemotherapy treated patients were generated using ROC plotter portal (http://www.rocplot.org/site/index) ((*68*). Gene correlation plots were obtained using cancer tool (CANCERTOOL (cicbiogune.es) using different breast cancer datasets.

### Statistical analysis

Data are presented as mean ±SD. To compare between experimental groups, we used Student’s t-test for experiments. The reported p-values associated with dataset analysis (TCGA or TCPA) are generated using online portals. p-value <0.05 is considered significant.

## Acknowledgements

This research work was supported by grants from Start-up grant (SRG/2021/001502) from Science and Engineering Research Board (SERB), Basic and Translational Research in Cancer grant (No.1/3(7)/2020/TMC/R&D-II/8823 Dt.30.07.2021), Capacity Building and Development of Novel and Cutting-edge Research Activities (No.1/3(4)/2021/TMC/R&D-II/15063 Dt.15.12.2021) from Department of Atomic Energy (DAE), Government of India.

“MKJ was supported by Ramanujan Fellowship (SB/S2/RJN-049/2018) awarded by Science and Engineering Research Board (SERB), Department of Science and Technology, Government of India”.

## Author contributions

NC: designed the study, performed experiments, analyzed the data, and written parts of the manuscript; BSC, SP and SM: designed and performed the experiments, analyzed data and written parts of manuscript; AS, SK, and SM: designed and performed experiments, analyzed data; EJ, SP, SM and VVK performed staining of human breast tissue sections, imaging and analysis; SS and SR: analyzed bioinformatics datasets and written parts of manuscript; MKJ: designed bioinformatics analysis methodology, reviewed and edited manuscript; SND: critically reviewed and edited manuscript; NV: designed the conception, developed experimental methods and models, acquired funding, performed and analyzed experimental and in-silico data, supervised and compiled the study, and prepared the manuscript. All authors read and approved the final manuscript.

