## Supplementary material for "GPX4-VIM equates a proliferating DTP state in TNBC subtypes with converged vulnerabilities to autophagy and glutathione inhibition"

***Running title***: GPX4-VIM equates a targetable DTPP state in TNBC.

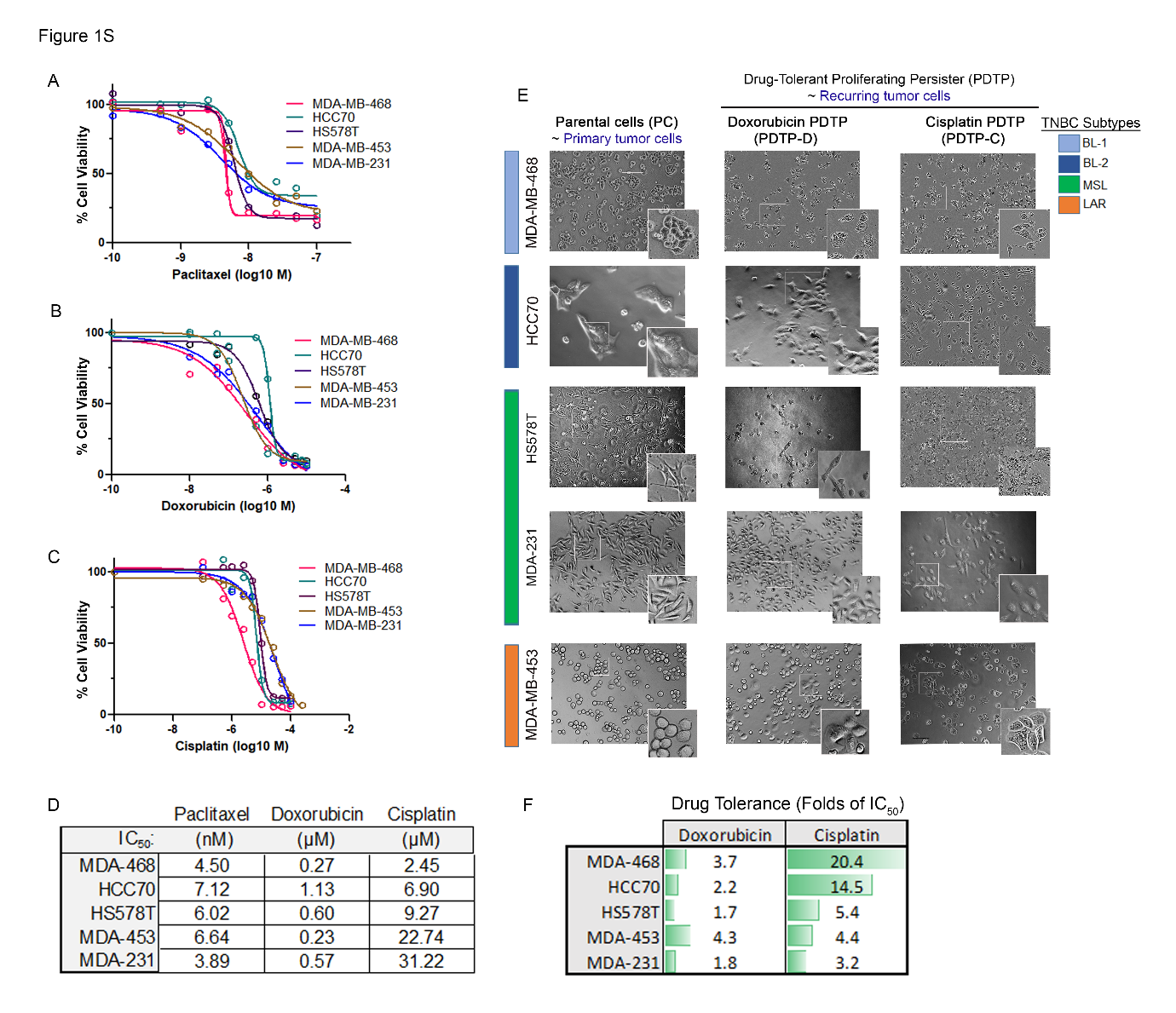

**Figure S1: Development of chemotherapy-tolerant TNBC persister cells from four molecular subtypes using different chemotherapeutic agents.**

(A, B, C) Dose response curve representing the percent cell viability of different subtypes of TNBC cells after 72-hour treatment with increasing concentration of Paclitaxel, Doxorubicin and Cisplatin respectively (D) Estimated IC_50_ values of drugs in different TNBC cell lines were listed in table (E) Bright field microscopy images and their zoomed inserts showing morphological differences between the different subtypes of TNBC PC and chemotherapy-tolerant proliferating persister cells. (F) Drug tolerance to doxorubicin and cisplatin represented as folds of folds of IC_50_ concentration of drugs in TNBC cell lines.

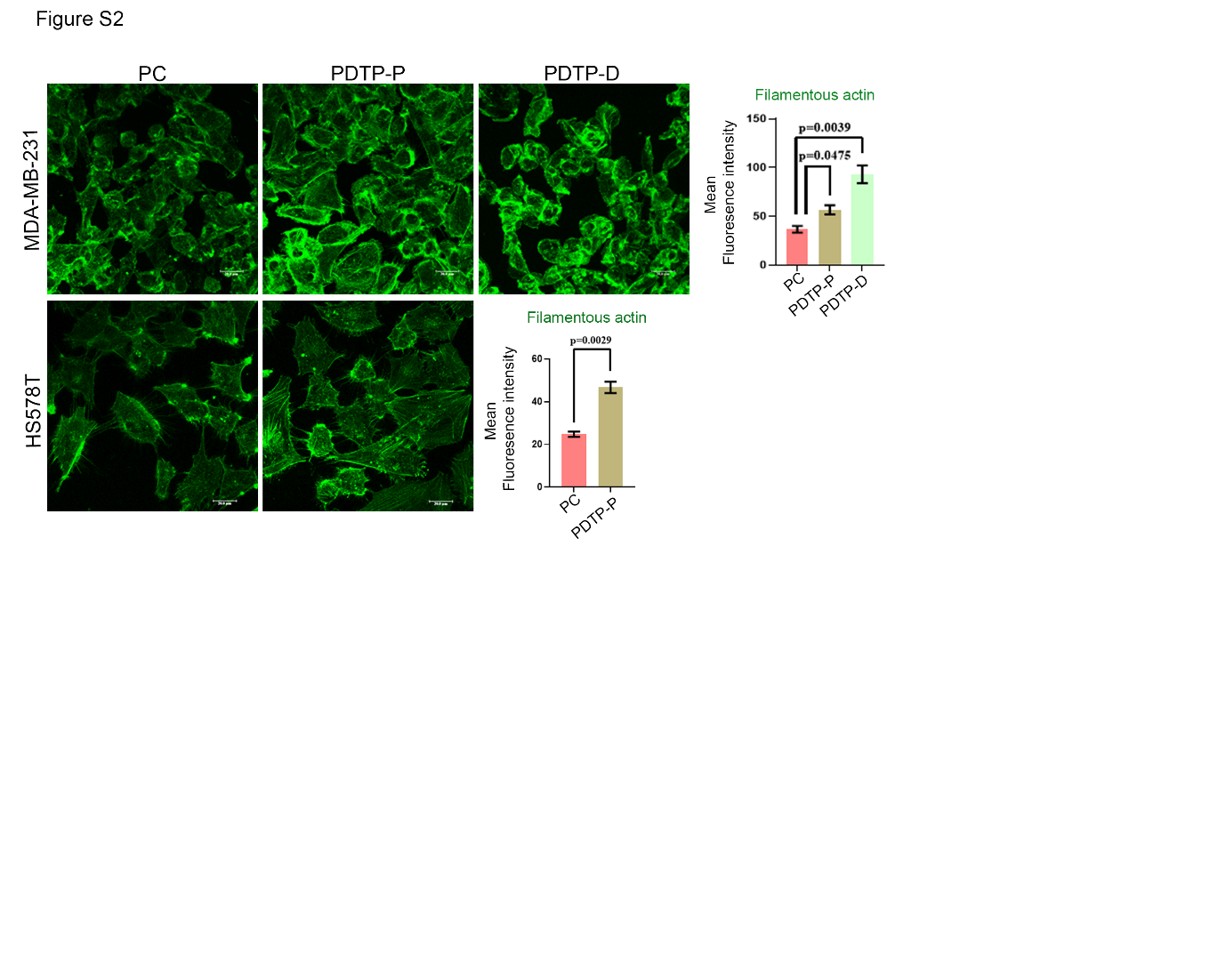

**Figure S2:** Confocal images showing filamentous actin staining and mean fluorescence intensity represented as bar graph in TNBC PC and respective PDTP cells.

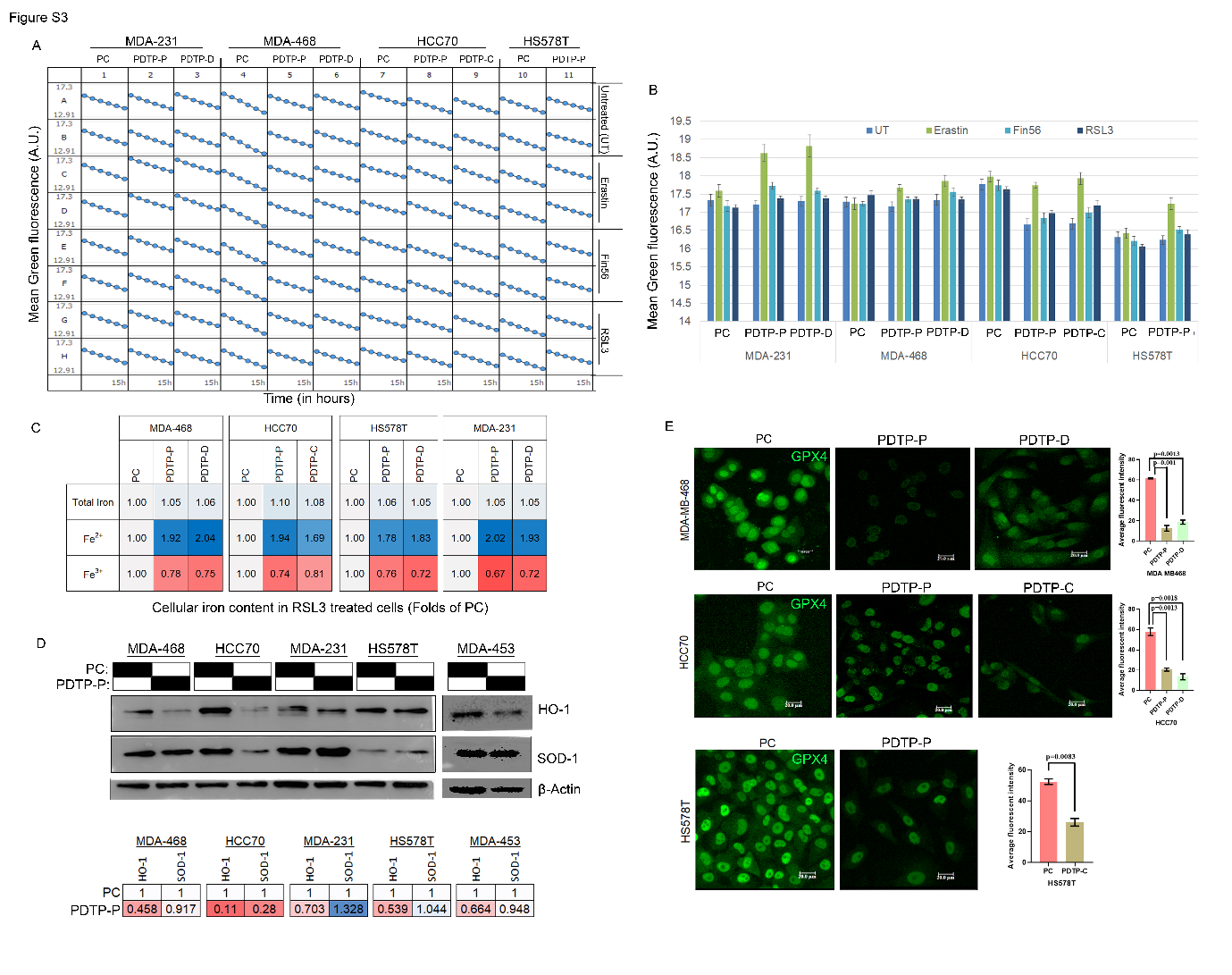
**Figure S3:** (A) Lipid peroxidation assessed using BODIPY 581/591 C11 lipid probes and mean green fluorescence was estimated at each time point time till 15h in different TNBC PC and PDTP cells upon treatment with different ferroptosis inducers as indicted in the representative microplate graphs. (B) Lipid peroxidation shown as mean±SD green fluorescence intensity at 4 hours of treatment with indicated ferroptosis inducers in PC and PDTP TNBC cell lines (C) Levels of HO-1 and SOD-1 protein expression in TNBC PC and PDTP-P cells shown by western blotting and band intensity was quantified by ImageJ and normalised to housekeeping β-actin, data shown as heatmap of expression values folds of PC. (D) Immunofluorescence images representing GPX4 staining and bar graph representing average fluorescent intensity of GPX4 staining in PC and PDTP cells of different TNBC cell lines.

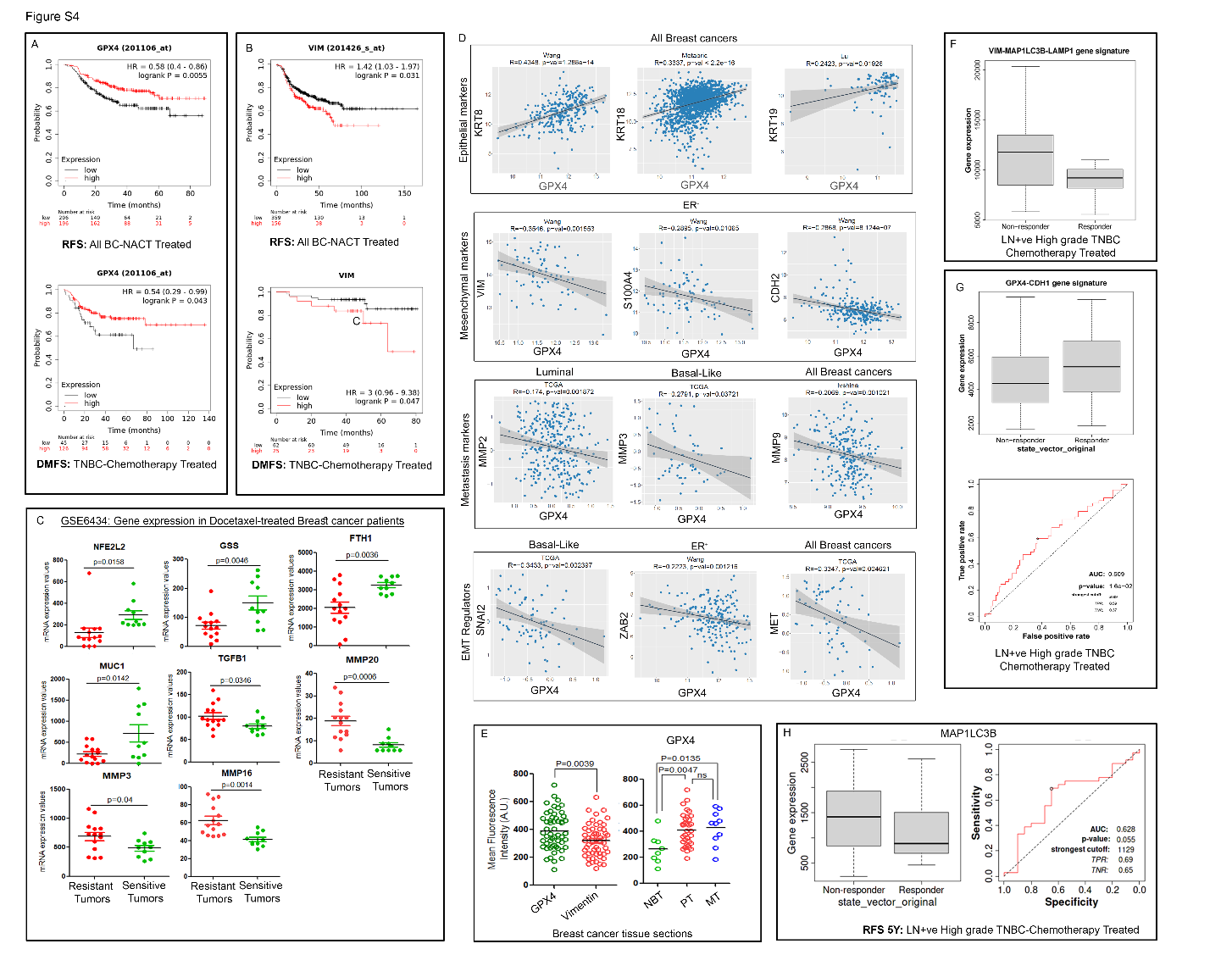
 **Figure S4**: **Association of GPX4, VIM and MAP1LC3B gene expressions with EMT, patient survival and response to chemotherapy in breast cancer.** (A) Kaplan-Meier Plot representing relapse free survival (RFS) and distant metastasis free survival (DMFS) in all breast cancer patients treated with neoadjuvant chemotherapy and TNBC patients treated with chemotherapy, respectively with high or low gene expression of GPX4 and (B) vimentin. (C) Expression of Ferroptosis genes in gene expression Affymetrix dataset (GSE6434) using GEOR: NFE2L2, GSS and FTH1; EMT genes: MUC1 and TGFB1; Metastasis genes: MMP20, MMP3, and MMP16 in breast cancer patients’ groups sensitive or resistant to docetaxel. (D) Correlation analysis of GPX4 gene with epithelial markers, mesenchymal markers, metastasis markers and EMT regulator genes in different breast cancer types taken from TCGA, metabric, Lu, Wang and Ivshina datasets. (E) Dot plots showing the Mean fluorescence intensities of GPX4 protein expression (right) in human breast tissue sections categorised under normal breast (NBT), primary tumor (PT) and lymph node metastatic tumors (MT) and for relative expression of GPX4 and VIM proteins in all the breast tissue sections (left) after performing immunofluorescence co-staining and quantitative analysis of paraffinized tissue sections. Each dot indicates a human breast tissue section analysed. (F) Box plot showing differential expression of MAP1LC2B gene between chemotherapy responder and non-responder TNBC patients with LN+ve high grade tumors, and its ROC curve analysis to predict RFS in these patients. (G) Expression of VIM-MAP1LC3B-LAMP1 tri-gene signature in chemotherapy responder vs non-responder TNBC patients’ groups with LN+ve high grade tumors. (H) Expression of GPX4-CDH1 gene signatures in chemotherapy responder vs non-responder TNBC patients’ groups with LN+ve high grade tumors and ROC curve analysis to predict survival.

| Sr. no. | Gene name | Primer sequence |
| --- | --- | --- |
| 1 | GPX4 | Forward- 5’GCCTTCCCGTGTAACCAGT3’ |
|  |  | Reverse- 5’GCGAACTCTTTGATCTCTTCGT3’ |
| 2 | GCLC | Forward- 5’AGACATTGATTGTCGCTG3’ |
|  |  | Reverse- 5’TGGTCAGACTCATTAGCA3’ |
| 3 | HO-1 | Forward- 5’ATGACACCAAGGACCAGA3’ |
|  |  | Reverse- 5’GTGTAAGGACCCATCGGA3’ |
| 4 | GSS | Forward- 5’TGCTAAAGCCCCAGAGAG3’ |
|  |  | Reverse- 5’AGCAGGCAATTCTCAAAAGG3’ |
| 5 | VIM | Forward- 5’ ACCGCTTTGCCAACTACAT3’ |
|  |  | Reverse- 5’ TTGTCCCGCTCCACCTC3’ |
| 6 | CDH1 | Forward- 5’GGAAGTCAGTTCAGACTCCAGCC3’ |
|  |  | Reverse- 5’AGGCCTTTTGACTGTAATCACACC3’ |
| 7 | Actin | Forward- 5’CATGAAGATCAAGATCATCGCC3’ |
|  |  | Reverse-ACATCTGCTGGAAGGTGGACA3’ |
| 8 | GAPDH | Forward-5’ CAATGACCCCTTCATTGACC3’ |
|  |  | Reverse-5’TTGATTTTGGAGGGATCTCG3’ |

Table S4. List of forward and reverse primer with their DNA sequences for QPCR amplification used in the study.
